# Spatial Glyco-Codes Define Human Liver Pathology and Progression

**DOI:** 10.64898/2026.07.08.737217

**Authors:** Xiaolong Tian, Anthony Fung, Xingbo Shang, Dingyao Zhang, Binfan Chen, Lei Zhang, Keyi Li, Mei Zhong, Yifan Deng, Mingyu Yang, Yao Lu, Bo Tao, Fu Gao, Alev Baysoy, Xiao Lin, Aleksandra Ivovic, Sidi Chen, Frank Li, Mina L. Xu, Xuchen Zhang, Mark Gerstein, Xiaoyong Yang, Chen Liu, Rong Fan

## Abstract

Glycosylation is a fundamental process regulating cellular function, tissue organization, and disease progression. However, comprehensive glycan profiling at single-cell spatial resolution remains largely inaccessible, particularly in clinical archival tissues. Here we develop spatial-GPT, a multimodal platform for simultaneous profiling of glycans, proteins, and/or transcripts in archival formalin-fixed paraffin-embedded (FFPE) tissues. Using a panel of 30 DNA-encoded lectins recognizing major mammalian glycan motifs and structural classes, sequencing-based spatial-GPT (DBiT-GPT) mapped the spatial glycome, proteome, and transcriptome across 16 human liver specimens encompassing steatosis, fibrosis, cirrhosis, and hepatocellular carcinoma (HCC), leading to identification of spatial glyco-codes – combinatorial glycan states associated with distinct cellular identities, tissue features, and pathological processes. Unexpectedly, glyco-codes alone were sufficient to resolve major cell types, disease states, and HCC subtypes, revealing a previously unappreciated level of biological information encoded within the tissue glycome. Spatial glycomics uncovered tumor-like glyco-codes in premalignant regions, suggesting that glycan reprogramming may precede overt malignant transformation. Using imaging-based single-cell spatial glycan-protein profiling (CODEX-GP), we track glyco-codes across the whole-tissue architecture of 3 representative HCC samples. We further examined the glyco-codes across more than 300 patient specimens and quantified cell-type- and disease-specific glyco-codes as well as glycan-defined immune-evasion, T-cell-exhaustion, and steato-fibrotic niches. Together, these findings establish spatial glyco-codes as a previously unrecognized layer of tissue organization that encodes cellular identity, tissue function, and disease progression. The ability of glyco-codes to distinguish major liver pathologies across independent patient cohorts further highlights their potential as a new class of molecular histopathology biomarkers.

## INTRODUCTION

Glycans are structurally diverse carbohydrate chains covalently attached to proteins and lipids that regulate the folding, trafficking, and molecular interactions of these biomolecules,^1^ thereby regulating cellular function, receptor signaling, cell–cell communication, and cell-matrix interactions. Accumulating evidence has further demonstrated the critical roles of glycosylation in immune recognition,^2^ host-pathogen interactions,^3^ and essentially every stage of tumor development.^4^ More recently, glycoRNAs have been increasingly recognized and have been shown to play critical roles in neutrophil recruitment.^5^ In order to understand the function of glycans in the tissue context, it is becoming increasingly important to characterize all major glycans and their modifications directly the tissue context.

scGlycan-seq was developed to detect glycans at single cell level and subsequently integrated with transcriptomic readouts to reveal cellular heterogeneity and the biological roles of glycans across multicellular systems.^6^ Mass spectrometry imaging is currently the mainstay technique for glycan profiling with spatial resolution. A representative platform is matrix-assisted laser desorption/ionization mass spectrometry imaging (MALDI-MSI), which is relatively unbiased and particularly well established for spatial mapping of released N-glycans. In FFPE tissues, MALDI-MSI can routinely detect 30 N-glycan species, distinguish tumor from non-tumor.^7^ Although MALDI-MSI has enabled spatially resolved glycan mapping, an inherent limitation is that many released glycans display identical or highly similar m/z values, restricting unambiguous structural and linkage-level interpretation. In addition, released-glycan MALDI-MSI does not directly preserve glycoprotein-carrier information and is not readily combined with same-section RNA or high-plex protein-marker profiling, thereby limiting multimodal integration and interpretation of glycan patterns across distinct cellular populations.

Here, we developed a totally different approach, utilizing a panel of 30 DNA-barcoded lectins to detect lectin-binding glycan motifs, further integrated for spatial co-profiling of Glycans, Proteins, and/or total RNA Transcripts (spatial-GPT) in human FFPE tissues. It comprises two complementary techniques: DBiT-GPT: a spatial sequencing approach built upon deterministic barcoding in tissue (DBiT) to realize glycan-included spatial multi-omics sequencing,^8^ and CODEX-GP: an optical imaging approach for single-cell spatial co-profiling of a panel of lectin-binding glycans and a panel of protein markers through Co-detection by indexing (CODEX).^9^ We applied DBiT-GPT to 16 human hepatocellular carcinoma (HCC) or its non-tumor background FFPE blocks that had been stored for 2-8 years. Surprisingly, glycans are extremely robust in FFPE tissues and successfully profiled across all samples including the ones stored over 8 years. Spatially distinct glycomic patterns (glyco-codes) were identified across normal and disease-associated spatial niches, defining glyco-codes, which were found to be also closely aligned with spatial transcriptomic architecture. Spatial co-sequenced proteins further facilitated canonical cell-type annotation. Using DBiT-GPT, we mapped glycome, proteome and transcriptome features across distinct cell types, resolved different HCC subtypes, and uncovered molecular programs associated with HCC progression. These findings revealed potentially promising therapeutic targets that are relatively conserved in HCC. In parallel, CODEX-GP enabled spatial co-imaging of glycans and proteins in these tissue samples at single-cell and subcellular resolution. Co-imaged proteins facilitated canonical cell-type annotation and enabled association of lectin-binding glycan motifs with protein-defined cell states and subcellular neighborhoods, revealing single-cell glycol-codes associated with steatotic hepatocytes, fibrosis, and distinct HCC subtypes. Further applying CODEX-GP to more than 300 patient specimens across a wide range of liver diseases quantified cell-type- and disease-specific glyco-codes as well as glycan-defined immune-evasion, T-cell-exhaustion, and steato-fibrotic niches. Together, these findings establish spatial glyco-codes as a previously unrecognized layer of tissue organization that encodes cellular identity, tissue function, and disease progression. The ability of glyco-codes to distinguish major liver pathologies across independent patient cohorts further highlights their potential as a new class of molecular histopathology biomarkers.

## RESULTS

### Rationale and development of DBiT-GPT

To enable sequencing-based spatial co-profiling of glycans, proteins, and transcripts in FFPE tissues, sections were first subjected to deparaffinization and crosslink reversal, followed by enzymatic in situ polyadenylation **(Figure 1A)**. This step expands RNA detection beyond intact polyadenylated mRNAs and allows the capture of diverse RNA species, including fragmented mRNAs and noncoding RNAs such as lncRNAs, miRNAs, and tRNAs.^10^ Next, tissue sections were stained with DNA-barcoded lectins to label both membrane-associated and intracellular glycans. Because tissue-bound lectins could potentially interact with glycosylated epitopes on subsequently applied antibodies, sections were briefly fixed and then blocked with a sugar mixture to saturate lectin-binding sites. High-plex protein targets were then profiled using a DNA-barcoded antibody cocktail.^11^ Reverse transcription (RT) was initiated in situ using a biotinylated oligo(dT) RT primer complementary to the polyadenylated transcripts as well as lectin- and antibody-derived DNA tags (LDTs and ADTs) bearing synthetic poly(A) tails. Spatial barcoding was subsequently performed using two orthogonal 50-channel microfluidic chips, generating a 2D grid of spatially barcoded tissue pixels. After imaging, tissues were lysed to recover spatially barcoded cDNA, LDTs, and ADTs, which were subsequently processed through template switching, library preparation, and sequencing **(Figure 1A)**.

**Figure 1.**
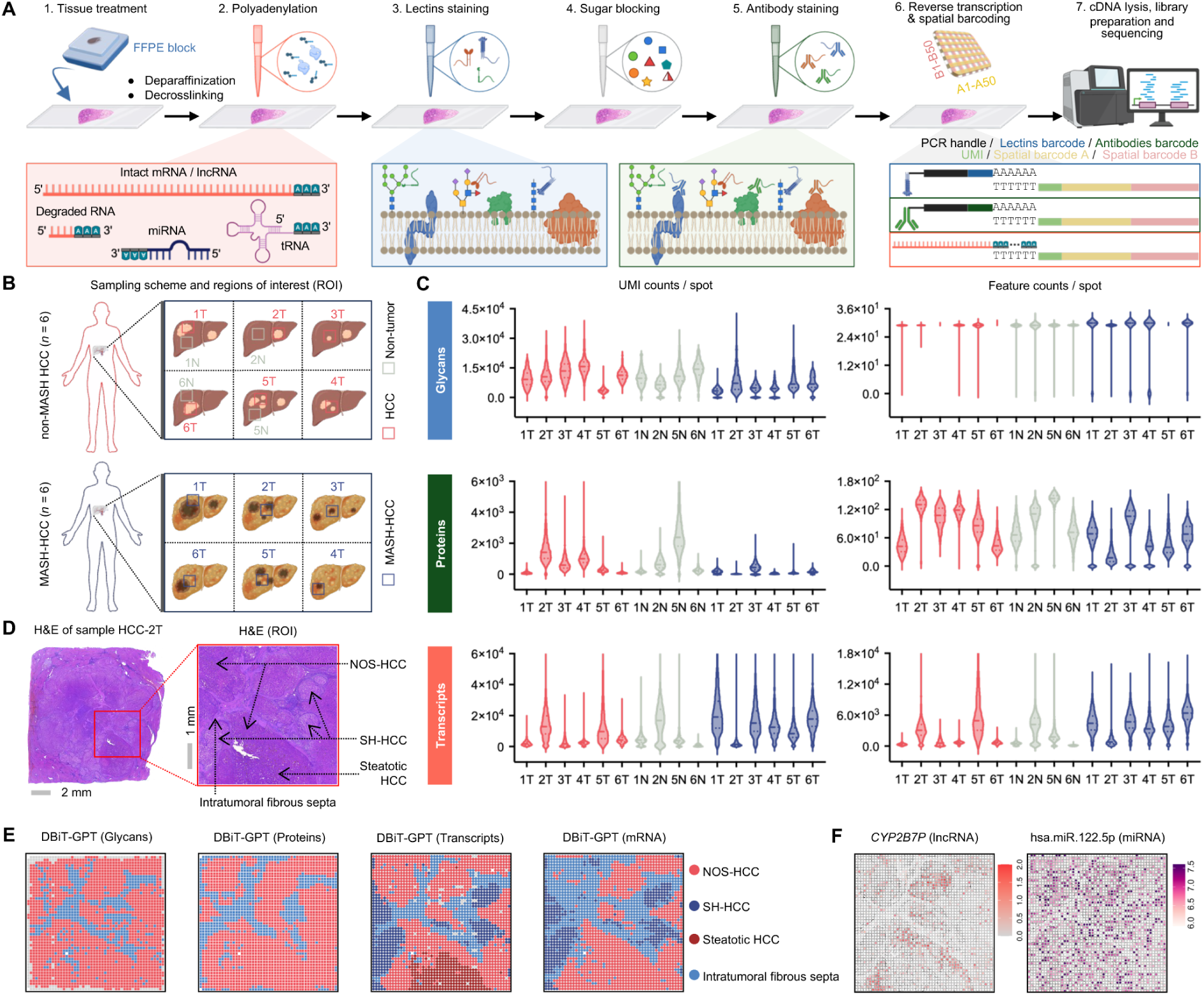
DBiT-GPT workflow and spatial mapping of glycans, proteins, and RNA transcripts across a variety of liver disease tissues. (A) Schematic workflow and chemical basis of DBiT-GPT. (B) Schematic representation of the HCC cohort and sequencing ROIs. (C) Distribution of UMI counts and detected feature counts per spot for glycans, proteins, and transcriptomes across HCC samples at 50 × 50 μm spatial resolution. (D-F) Spatial clustering results of DBiT-GPT in sample HCC-2T. (D) H&E staining of an adjacent section. (E) Unsupervised clustering based on spatial glycome, proteome, transcriptome, and mRNA-only transcriptome data, together with individual spatial plots of selected ncRNAs (F). See also Figures S1 and S2.

The lectin panel used in this study comprised 30 well-characterized lectins **(Table S2)** assessed by the National Center for Functional Glycomics (NCFG). This panel captures major mammalian glycan motifs, including fucose and Lewis-type structures, mannose-rich and core-fucosylated N-glycans, terminal galactose/LacNAc, GlcNAc/LacNAc-containing structures, GalNAc- and T/Tn antigen-related O-glycans, α2,3- and α2,6-sialylated structures, and branched or bisected complex N-glycans. In parallel, the human antibody Total-seq cocktail targets 154 unique cell-surface antigens **(Table S2)**, including principal lineage and functional markers. Together with transcriptome-wide total RNA profiling, these glycan, protein, and transcript readouts enable DBiT-GPT to spatially profile and integrate the glycome, proteome, and transcriptome all from a single FFPE tissue section.

### DBiT-GPT exhibits robust spatial multi-omics sequencing capability in archived clinical FFPE HCC samples

To evaluate the robustness of DBiT-GPT in archived clinical FFPE specimens, we applied the workflow to 16 HCC FFPE blocks: 6 tumor regions and 4 matched non-tumor background liver regions from 6 non-MASH HCC patients, and 6 tumor regions from another 6 metabolic dysfunction-associated steatohepatitis-associated HCC (MASH-HCC) patients **(Figure 1B)**. These blocks had been archived for 2 to 8 years **(Figure S1A)**.

As expected, sequencing results showed that transcript yield was associated with tissue quality including FFPE block storage duration. Six blocks with shorter storage durations showed more consistent transcript yields, with UMI counts distributed between ∼10,000 and 20,000 UMIs per pixel, whereas the 10 blocks archived for approximately 8 years showed lower but still robust transcript yields, ranging from ∼2,000 to 20,000 UMIs per pixel **(Figures 1C and S1B)**. In contrast, glycan-associated LDT yields remained robust in long-archived blocks, with most blocks archived for approximately 8 years yielding more than 10,000 LDT UMIs per pixel. LDT yields were generally lower in MASH-HCC samples, likely reflecting steatosis-associated tissue-context effects, although nearly all 30 LDT features were detected across most spots **(Figures 1C and S1B)**. Protein-associated ADT yields were consistent across samples, with most pixels detecting approximately 50–100 protein features **(Figures 1C and S1B)**. Together, these results indicate that DBiT-GPT is highly reliable even for ultra-long-term archival FFPE samples, while the variation of glycan yield actually reflects the biology of tissues themselves.

HCC-2T was a non-MASH HCC sample whose ROI contained multiple histological components, including not-otherwise-specified HCC (NOS-HCC), hepatitis C virus (HCV)-associated steatohepatitic HCC (SH-HCC), and regions of focally steatotic HCC **(Figures 1D and S1A)**. Unsupervised clustering of DBiT-GPT spatial glycome and proteome data accurately mapped the HCC tumor regions and the intratumoral fibrous septa separating tumor colonies, with highly concordant spatial patterns between the two modalities. These spatially aligned glycan and protein maps further enabled cell type–resolved co-localization analysis, in which pairwise Pearson correlations identified proteins spatially associated with representative cell type–associated lectin-binding glycans **(Figures S1E and S1F)**. Classical mRNA-based spatial clustering distinguished SH-HCC from NOS-HCC, but the upper-left NOS-HCC region partially clustered with fibrotic regions. While spatial total-transcriptome clustering provided finer resolution and precisely resolved all major histological morphologies in the ROI **(Figures 1E, S1C, and S1D)**. Extending beyond mRNA, noncoding RNAs were detected such as widely observed expression of hsa-miR-122, a previously reported diagnostic biomarker for HCC and non-alcoholic fatty liver disease (NAFLD),^12^ across the ROI **(Figure 1F)**. In addition, the lncRNA *CYP2B7P*, a shared target of hsa-miR-122 and another HCC-associated biomarker, hsa-miR-183-5p,^13^ showed selective enrichment in the NOS-HCC region **(Figure 1F)**. In summary, these data demonstrate that DBiT-GPT is effective in unraveling histological heterogeneity and complex molecular patterns in archived clinical FFPE tissue sections.

In another non-MASH HCC sample, HCC-3T **(Figure S2B)**, all three spatial modalities mapped tumor and non-tumor regions **(Figure S2A)**. Notably, the spatial glycome more precisely resolved focal SH-HCC-like and necrotic subregions, whereas the spatial proteome delineated scattered NOS-HCC and necrotic subregions **(Figures S2A and S2C)**. In non-tumor background sample of HCC-6N **(Figure S2D)**, where RNA degradation limited transcript profiling to only a small number of detected transcripts **(Figure S2E)**, the spatial glycome and proteome still accurately mapped non-tumor histological structures, including cirrhotic nodule hepatocytes (CH) and the fibrous septa connecting adjacent nodules **(Figures S2F and S2G)**. In conclusion, these results highlight the concordance and complementarity of DBiT-GPT glycome, proteome, and transcriptome readouts, even in archived FFPE tissues with variable RNA quality.

### DBiT-GPT maps progression of NOS-HCC

Liver cancer represents a growing global health burden, and hepatitis B virus (HBV) and HCV infections remain leading causes **(Figure 2A)**.^14^ Given that clinical monitoring of liver disease progression toward liver cancer primarily relies on glycoprotein- and glycosylation-based biomarkers such as alpha-fetoprotein (AFP) and AFP-L3,^15^ we hypothesized spatial glycome may be sufficient to differentiate these pathological conditions. Thus, DBiT-GPT was applied to dissect the spatial molecular mechanisms underlying several major HCC pathologies including virus-associated hepatocellular carcinoma and NOS-HCC progression. Sample of HCC-5T was derived from an HCV-infected patient with NOS-HCC **(Figures 2B, S1A, and S3A)**. Both spatial glycome and transcriptome data mapped the major histopathological regions within the ROI, including cirrhotic hepatocytes with steatosis, fibrous septa, and early-stage NOS-HCC **(Figures 2C**–**2F and S3A–S3C)**. Based on differentially expressed glycans of each clusters, we observed and defined three glyco-codes: glyco-code 1, consisting of MPL-, Con A-, and DBA-binding glycans; glyco-code 2, consisting of GSL I-, GSL I-B4-, LCA-, PHA-E-, PHA-L-, PSA-, UEA I-, VVL-, and WFA-binding glycans; and glyco-code 3, consisting of DSL-, AAL-, LEL-, Con A-, and RCA I-binding glycans **(Figures 2F and S5A–S5C)**. These glyco-codes showed strong spatial concordance with cirrhotic hepatocytes with steatosis, fibrous septa, and NOS-HCC, respectively **(Figure 2G)**, and were dominantly enriched in their corresponding glycan clusters **(Figure 2H)**.

**Figure 2.**
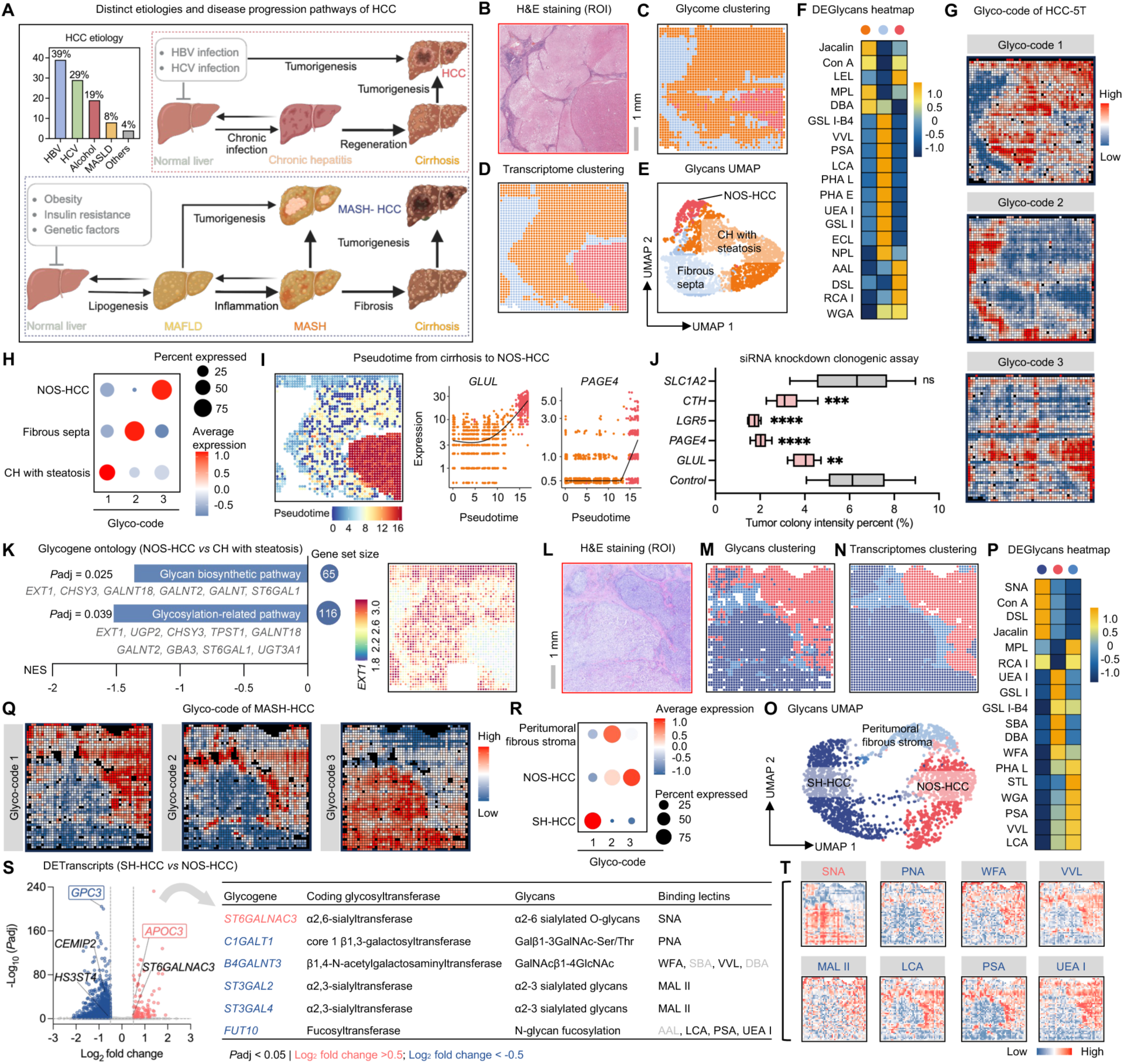
DBiT-GPT maps progression of NOS-HCC and resolves tumor heterogeneity in MASH-HCC. (A) Top left: etiologies of distinct HCC subtypes, adapted from the World Health Organization (WHO). Top right: disease progression pathway of HBV/HCV-associated HCC. Bottom: disease progression pathway of MASH-driven HCC. (B–K) All data shown in panels B–K are from an HCV-associated NOS-HCC sample (sample ID: HCC-5T). (B) ROI from H&E staining of an adjacent section. (C–D) Spatial maps of unsupervised clustering based on glycans (C) and transcriptomes (D) from the same HCC-5T section. (E) UMAP of glycan clusters identified in the HCC-5T. (F) Heatmap of the top five differentially expressed glycans defining each cluster. (G–H) Definition and spatial expression patterns of glyco-codes derived from the HCC-5T spatial glycome. (G) Spatial maps of the HCC-5T glyco-codes. (H) Dot plot summarizing the expression of each glyco-code across spatial glycan clusters. (I) Left: pseudotime analysis of the HCC-5T spatial transcriptome generated by DBiT-GPT, revealing a pseudotemporal continuum from CH with steatosis to an early stage of NOS-HCC. Middle and right: representative transcripts dynamically changing along pseudotime. (J) Clonogenic assay following siRNA knockdown of top transcripts identified in NOS-HCC cluster, including *GLUL* and *PAGE4*, which were also among the top transcripts dynamically changing along pseudotime. (K) Glycogene ontology analysis comparing clusters of NOS-HCC and CH with steatosis. Left: two significantly enriched pathways (*P*adj < 0.05). Right: representative spatial expression of glycogene *EXT1*. (L–T) All data shown in panels L–T are from a MASH-driven HCC sample (sample ID: MASH-HCC-1T). (L) ROI from H&E staining of an adjacent section. (M–N) Spatial maps of unsupervised clustering based on DBiT-GPT glycome (M) and transcriptome (N) from the same MASH-HCC-1T section. (O) UMAP of glycan clusters identified in the MASH-HCC-1T. (P) Heatmap of the top seven differentially expressed glycans defining each cluster. (Q–R) Definition and spatial expression patterns of glyco-codes derived from the MASH-HCC-1T spatial glycome. (Q) Spatial maps of the MASH-HCC-1T glyco-codes. (R) Dot plot summarizing the expression of each glyco-code across spatial glycome clusters. (S) Left: differentially expressed transcripts between SH-HCC and NOS-HCC. Right: significant glycogenes identified from the differentially expressed transcripts and their functions. (T) Spatial plots of individual lectin-binding glycans corresponding to the differentially expressed transcripts shown in panel (S). See also Figures S3, S4, S5, and S6.

Notably, the NOS-HCC-associated glyco-code was also elevated in cirrhotic regions adjacent to the tumor **(Figure 2G)**, suggesting that glycosylation patterns characteristic of NOS-HCC may already be present in premalignant tissue. Tandem transcriptomic sequencing confirms several glycosyltransferase genes in these regions were reduced, potentially impairing downstream proteomic function. Together, these results suggest that even morphologically non-malignant histology may already be reprogramming its metabolic profile towards malignancy, and glyco-codes may offer an early biomarker.

To define the programs associated with this progression, we performed transcriptional pseudotime analysis across cirrhotic hepatocytes - a classical pathway of hepatocarcinogenesis **(Figure 2A)**^16^, with steatosis and NOS-HCC **(Figures 2I and S3D–S3E)**. Along this trajectory, runt-related transcription factor 1 (*RUNX1)*, a transcription factor implicated in tumor progression and cellular plasticity, was progressively increased,^17^ whereas apolipoprotein C3 (*APOC3)*, a gene involved in triglyceride-rich lipoprotein metabolism, was decreased **(Figures S3E)**.^18^ glutamate-ammonia ligase (*GLUL*), a canonical zone 3/pericentral hepatocyte marker, was markedly elevated **(Figures 2I and S3G)**, suggesting a shift toward a pericentral-like tumor program.^19^ Further, siRNA-mediated knockdown of NOS-HCC-enriched *GLUL*, *PAGE4*, *LGR5*, and *CTH* significantly reduced Huh7 colony formation **(Figures 2J–2K and S3E–S3F)**, suggesting potential therapeutic vulnerabilities in HCC.

Unexpectedly, exostosin glycosyltransferase 1 *(EXT1)*, a canonical glycogene^20^ was reduced in NOS-HCC **(Figures 2K and S3E)**. Ontology analysis of differentially expressed glycogenes^21^ further showed that the only two significantly altered GlycoEnzyme pathway were both downregulated in NOS-HCC **(Figure 2K)**. These findings indicate that aberrant glycosylation gene regulation arise earlier during premalignant liver disease progression, even though NOS-HCC displays enhanced glycolytic activity **(Figures S3H)**.

### DBiT-GPT resolves tumor heterogeneity in MASH-HCC

Among major liver cancer etiologies **(Figure 2A)**, Metabolic Dysfunction-Associated Steatotic Liver Disease (MASLD) is uniquely increasing in both age-standardized incidence and mortality,^14^ highlighting the need to better understand the mechanisms of MASH-HCC. We therefore applied DBiT-GPT to a MASH-driven HCC sample, MASH-HCC-1T **(Figures 2L, S1A, and S4A)**. Sequencing results revealed that SH-HCC and NOS-HCC regions displayed both distinct glycomic and transcriptomic patterns **(Figures 2M–2O, and S4B–S4C)**. Based on differentially expressed glycans of each clusters, we identified three glyco-codes in this sample: glyco-code 1, marked by SNA-, Con A-, DSL-, MPL-, and RCA I-binding glycans; glyco-code 2, marked by STL-, PSA-, PNA-, VVL-, PHA-L-, and LCA-binding glycans; and glyco-code 3, marked by UEA I-, SBA-, DBA-, GSL I-, GSL I-B4-, and WFA-binding glycans **(Figures 2P and S6A–S6C)**. These glyco-codes were selectively enriched in SH-HCC, peritumoral fibrous stroma, and NOS-HCC, respectively **(Figures 2Q–2R)**.

As expected, DBiT-GPT transcriptome results showed that *APOC3* and glypican 3 (*GPC3*), the latter being a canonical HCC biomarker,^22^ were predominantly expressed in SH-HCC and NOS-HCC regions, respectively **(Figures 2S and S4D)**. Our unique spatial transcriptome sequencing allows for genome-wide single nucleotide variant (SNV) profiling and SNV-based clone analysis further revealed distinct transcriptome-inferred somatic variant patterns between these two tumor components, supporting their divergent genetic backgrounds **(Figure S4E)**. However, glycogene ontology did not identify significantly different GlycoEnzyme pathways between SH-HCC and NOS-HCC (**Figures S4F–S4G**). We further compared the differentially expressed genes between these two components with a curated set of 403 human glycoEnzymes coding genes, and the result showed distinct glycogene expression programs in SH-HCC and NOS-HCC **(Figure 2S)**. For example, *ST6GALNAC3* was significantly enriched in SH-HCC **(Figure 2S and S4D)**. *ST6GALNAC3* encodes an α2,6-sialyltransferase involved in the biosynthesis of α2,6-sialylated O-glycans, a glycan class that can be recognized by the lectin of SNA,^23^ consistent with the enriched SNA-derived LDT signal in SH-HCC region **(Figures 2S**–**2T and S4H)**. Conversely, several glycogenes significantly enriched in NOS-HCC, including *C1GALT1*, *B4GALNT3*, *ST3GAL2*, *ST3GAL4*, and *FUT10*, are associated with glycosylation motifs recognized by lectins such as PNA, WFA, VVL, MAL II, LCA, PSA, and UEA I, whose LDTs signals were significantly higher in NOS-HCC than in SH-HCC **(Figures 2S**–**2T and S4H–4I)**.

Taken together, DBiT-GPT spatial multi-omic profiling revealed glycome and transcriptome spatial heterogeneity within MASH-HCC and connected distinct glycan phenotypes to their underlying glycogene regulatory programs pixel-by-pixel.

### DBiT-GPT decodes the spatial architecture of glycogen-rich clear cell HCC (HCC-CC)

In addition to SH-HCC, the HCC-CC represents another distinct histological variant characterized by glycogen-rich clear cytoplasm **(Figure 3A)**, which can partially overlap morphologically with SH-HCC due to admixed lipid droplets.^24^ Although associated with a more favorable prognosis than conventional HCC, its molecular features remain poorly defined **(Figure 3A)**.^24^ We next applied DBiT-GPT to spatially resolve the HCC-CC architecture. In the sample of HCC-6T **(Figures 3B, S1A, and S7A)**, unsupervised spatial clustering based on glycome, proteome, and transcriptome showed highly concordant anatomical organization across the three molecular modalities, accurately delineating HCC-CC, neighboring cirrhotic hepatocytes, and stromal septal regions **(Figures 3D**–**3F, S7B–S7C, and S7E–S7F)**. Specifically, glycan profiling identified two spatially segregated spatial glyco-codes associated with HCC-CC and zone 3-like cirrhotic hepatocytes, respectively **(Figures 3G**–**3H)**. Glyco-code 1, marked by RCA I-, LTL-, VVL-, WGA-, GSL II-, and LCA-binding glycans, and glyco-code 2, marked by PHA-L-, STL-, LEL-, and DSL-binding glycans **(Figures 3F and S8A–S8B)**. Among them, RCA I- and PHA-L-binding glycans closely matched their corresponding histopathological compartments **(Figures S8A–S8B)**. In addition, Con A, which recognizes α-mannosyl and α-glucosyl residues and is compatible with glycogen-associated carbohydrate-rich regions **(Figure S7D)**, showed broad enrichment across the ROI, consistent with the strong Periodic acid–Schiff (PAS) signal observed in this clear cell lesion **(Figure 3C)**.

**Figure 3.**
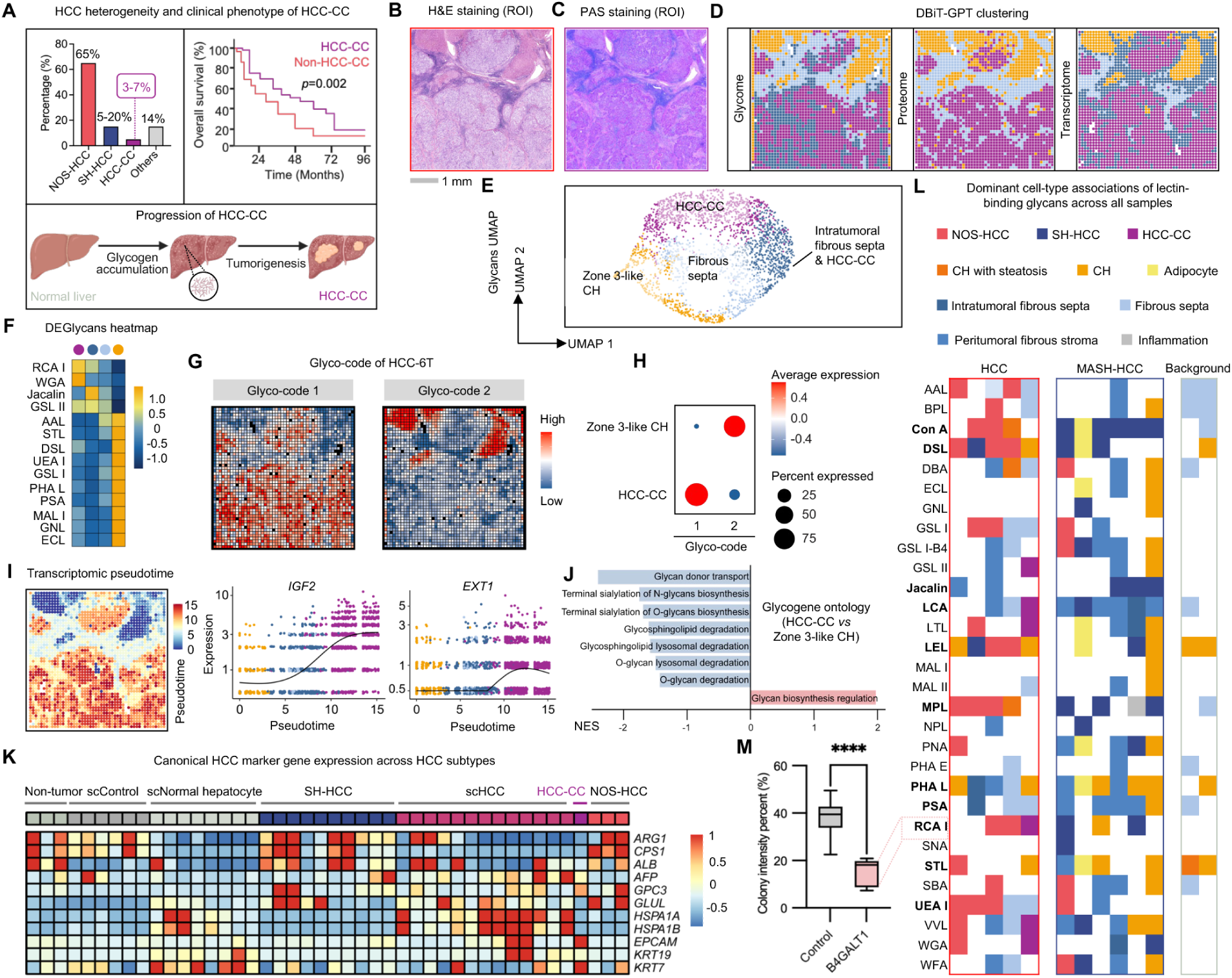
DBiT-GPT decodes the spatial architecture of glycogen-rich clear cell HCC. (A) HCC heterogeneity and clinical phenotype of HCC-CC. (B) ROI from the H&E staining of an adjacent section. (C) PAS staining of an adjacent section (corresponding ROI), highlighting glycogen accumulation, a characteristic feature of HCC-CC. (D) Spatial maps of unsupervised clustering of the same HCC-CC section. From left to right: glycome-, proteome-, and transcriptome-based clustering. (E) UMAP visualization of glycans clusters identified in the HCC-CC sample. (F) Heatmap of the top five differentially expressed glycans defining each cluster. (G) Definition and expression patterns of glyco-codes derived from DBiT-GPT. (H) Dot plot summarizing the expression patterns of each glyco-code across spatial glycan clusters. (I) Left: pseudotime analysis of the spatial transcriptome generated by DBiT-GPT, revealing a pseudotemporal continuum from Zone 3-like CH to HCC-CC. Right: representative transcripts dynamically changed along pseudotime. (J) Glycogene ontology analysis comparing HCC-CC and Zone 3-like CH. (K) Integrated heatmap of canonical HCC marker genes across HCC subtypes, generated by integrating DBiT-GPT datasets with published single-cell and single-nucleus RNA-seq datasets to reveal subtype-specific expression heterogeneity. (L) Dominant cell-type associations of lectin-binding glycans across all samples. Each tile indicates the most enriched cell type associated with a given LDT in each sample, grouped by non-MASH HCC (HCC), MASH-HCC, and background liver blocks. (M) Clonogenic assay showing reduced colony formation after siRNA-mediated knockdown of *B4GALT1*, which encodes a β1,4-galactosyltransferase mediating RCA I–binding glycan biosynthesis. See also Figures S7, S8, S9, S10, S11 and S12.

At the protein level, HCC-CC exhibited selective upregulation of CD63,^25^ LAMP-1,^26^ and CD39,^27^ revealing a distinct endosomal–lysosomal and vesicular trafficking program coupled with CD39-associated purinergic immunosuppressive signaling **(Figure S7E)**. To further interrogate the transition from adjacent cirrhotic hepatocytes to HCC-CC, we performed pseudotime analysis of the spatial transcriptomics data. The inferred trajectory revealed a continuum from zone 3-like cirrhotic hepatocytes toward HCC-CC **(Figures 3I and S7I),** indicating zone 3 cells may act as the cell of origin in HCC-CC pathogenesis. Along this axis, *IGF2* and *HFM1* progressively increased **(Figures 3I and S7I)**, consistent with activation of tumor growth^28^ and genome instability-associated programs,^29^ whereas canonical hepatocyte functional genes, including *ALB*, *HAMP*, *HRG*, *HP*, *HPR*, and *FGB*, gradually decreased **(Figure S7H)**. Similarly, LEL- and DSL-binding glycans progressively decreased from cirrhotic hepatocytes toward HCC-CC, paralleling the decline in mature hepatocyte functional genes **(Figure S8B)**.

We then examined glycogene programs to define the regulatory basis of this altered glycan landscape. Notably, *EXT1*, a glycosyltransferase involved in heparan sulfate biosynthesis, showed minimal expression in NOS-HCC but was enriched in cirrhotic hepatocytes in HCC-5T **(Figures 2K and S3E)**. This pattern contrasted with the HCC-CC sample, in which *EXT1* was preferentially elevated in the tumor region **(Figure 3I)**. Consistent with this distinction, glycogene ontology analysis revealed increased overall glycan biosynthesis regulation in HCC-CC, whereas adjacent zone 3-like cirrhotic hepatocytes displayed stronger enrichment of terminal sialylation biosynthesis and glycan donor transport pathways **(Figure 3J)**. This enrichment may reflect an increased capacity of cirrhotic hepatocytes to supply activated sugar donors, potentially supporting the broader glycan biosynthetic remodeling observed in HCC-CC. In parallel, QIAGEN Ingenuity Pathway Analysis (IPA) pathway analysis revealed enrichment of carbohydrate metabolism and glycosylation-related pathways in HCC-CC **(Figure S7J)**. Together, these findings reveal a spatial glycan expression and regulatory landscape in HCC-CC that is markedly distinct from conventional HCC.

Finally, to place HCC-CC within the broader landscape of HCC heterogeneity, we integrated spatial transcriptomes from all samples in this study with published single-cell^30^ and single-nucleus^31^ RNA-seq datasets. This cross-dataset analysis revealed substantial heterogeneity in canonical HCC marker expression across tumor subtypes. Overall, *ARG1*, *CPS1*, *ALB*, and *GLUL* were relatively enriched in NOS-HCC and SH-HCC but were largely absent in HCC-CC **(Figure 3K)**. This distinct marker expression pattern may partly explain why this variant is often associated with a more favorable prognosis than conventional HCC.

### Pan-sample glycan profiling reveals conserved cell type–specific glycan signatures

To further determine whether shared glycan patterns exist across HCC subtypes and other tissue compartments, we integrated the spatial glycome profiles from 13 samples after excluding 1 ROI composed entirely of tumor tissue and 2 samples composed entirely of normal liver **(Figure 3L)**. For each sample, we assigned the spatial distribution of all 30 LDTs signals to their corresponding dominant cell type or pathological compartment, with non-dominant or undetected associations shown as blank **(Figure 3L)**. Among tumor compartments, Con A-binding glycans were broadly enriched in 5 of 6 SH-HCC samples, suggesting a recurrent glycan feature associated with the SH-HCC phenotype **(Figures 3L and S9–S12)**. In contrast, UEA I-binding glycans were selectively enriched in 4 NOS-HCC compartments **(Figures 3L and S9–S12)**. DSL- and MPL-binding glycans showed a more balanced distribution, with broad enrichment across both NOS-HCC and SH-HCC **(Figures 3L and S9–S12)**. RCA I-binding glycans, in contrast, emerged as a more broadly conserved HCC-associated glycan program, with enrichment across the three major HCC subtypes examined in this study **(Figures 3L and S9–S12)**. To functionally evaluate the relevance of this broad HCC-associated RCA I-binding glycan program, we performed siRNA-mediated knockdown of beta-1,4-galactosyltransferase 1 (*B4GALT1*), which encodes a β1,4-galactosyltransferase that contributes to the biosynthesis of terminal β-galactosylated structures^32^ recognized by RCA I.^33^ *B4GALT1* knockdown markedly reduced colony formation in Huh7 cells **(Figure 3M)**, supporting a functional role for this RCA I–B4GALT1-associated glycan axis in HCC growth.

Beyond tumor-associated glycan programs, we also identified glycan patterns linked to other pathological states. LEL-binding glycans were strongly associated with steatosis compartments, showing enrichment not only in adipocytes but also in cirrhotic hepatocytes with steatosis and SH-HCC **(Figures 3L and S9–S12)**. In contrast, PHA-L- and STL-binding glycans were preferentially enriched across multiple cirrhotic hepatocyte compartments **(Figures 3L and S9–S12)**, suggesting a conserved association with cirrhotic or chronically injured hepatocyte states.

We further observed highly conserved fibrosis-associated glycan programs. Jacalin-binding glycans were strongly associated with cancer-associated fibroblast (CAF)-enriched tumor-associated fibrotic stroma, including intratumoral fibrous septa and peritumoral fibrous stroma **(Figures 3L and S9–S12)**. LCA- and PSA-binding glycans showed similarly conserved but broader enrichment across fibrotic compartments **(Figures 3L and S9–S12)**, indicating that core fucosylated and mannose-rich glycan structures may represent common features of HCC-associated stromal remodeling.

Collectively, these DBiT-GPT lectin-based spatial glycome profiles revealed conserved glycan features across distinct liver disease states, establishing a spatial glyco-code framework for understanding molecular heterogeneity in liver diseases.

### Rationale and development of CODEX-GP

Complementary to sequencing-based DBiT-GPT, which links spatial glycomics to protein markers and underlying transcriptional programs but does not provide subcellular resolution for single-cell whole-slide mapping of glycan expression, we also developed CODEX-GP as an imaging-based platform for co-mapping of glycans and proteins. Built upon DNA-encoded lectins and the CODEX framework,^9^ CODEX-GP enables multiplexed co-imaging of diverse glycan motifs and high-plex protein expression profiling within the same tissue section **(Figure S14A)**.

Given that streptavidin-biotin conjugation strategy used for DNA-barcoded lectins in DBiT-GPT is non-covalent, its stability would be reduced during repeated dimethyl sulfoxide (DMSO) wash steps across CODEX cycles. Therefore, a click chemistry-based strategy^34^ that generates a covalent linkage enables the robust multiplexing of 30 DNA-barcoded lectins **(Table S3)** for CODEX-GP. Specifically, lectins were conjugated to 5′ azide-modified DNA barcodes via DBCO-mediated click chemistry through lectin surface amines. Since this chemistry consumes amine groups that could otherwise support further paraformaldehyde (PFA) fixation, we introduced a 3′ amine modification on the barcode as an additional fixation handle to stabilize tissue-bound lectins during repeated CODEX imaging cycles **(Figure S14A, Table S3)**. We first applied CODEX-GP to three representative HCC specimens previously profiled by DBiT-GPT, including NOS-HCC (HCC-5T), MASH-HCC (MASH-HCC-1T), and HCC-CC (HCC-6T). Notably, SNA, SBA, and UEA-I were imaged in the final glycan imaging cycle, and their imaging signals remained stable with clearly preserved spatial patterns after 11 CODEX imaging cycles, as observed in the raw imaging files **(Figures S15A–S15C)**. Following the same strategy used in DBiT-GPT, lectin staining was performed first, followed by sugar blocking to saturate the glycan-binding sites of tissue-bound lectins before subsequent antibody staining. Manually annotated images show that after blocking non-specific lectin binding, even within tumor regions, there is significant heterogeneity in lectin signal **(Figure 4A)**. In total, 23 of 30 lectins were efficiently conjugated with barcodes and generated clear spatial glycan imaging signals **(Figures S15)**. Antibodies targeting 27 HCC progression–associated proteins **(Table S3)** were conjugated with DNA barcodes **(Table S3)** using thiol–maleimide chemistry following the CODEX antibody conjugation protocol **(Figure S14A)**.^9^ The resulting DNA-barcoded antibodies were subsequently applied to the same tissue section for proteins staining.

**Figure 4.**
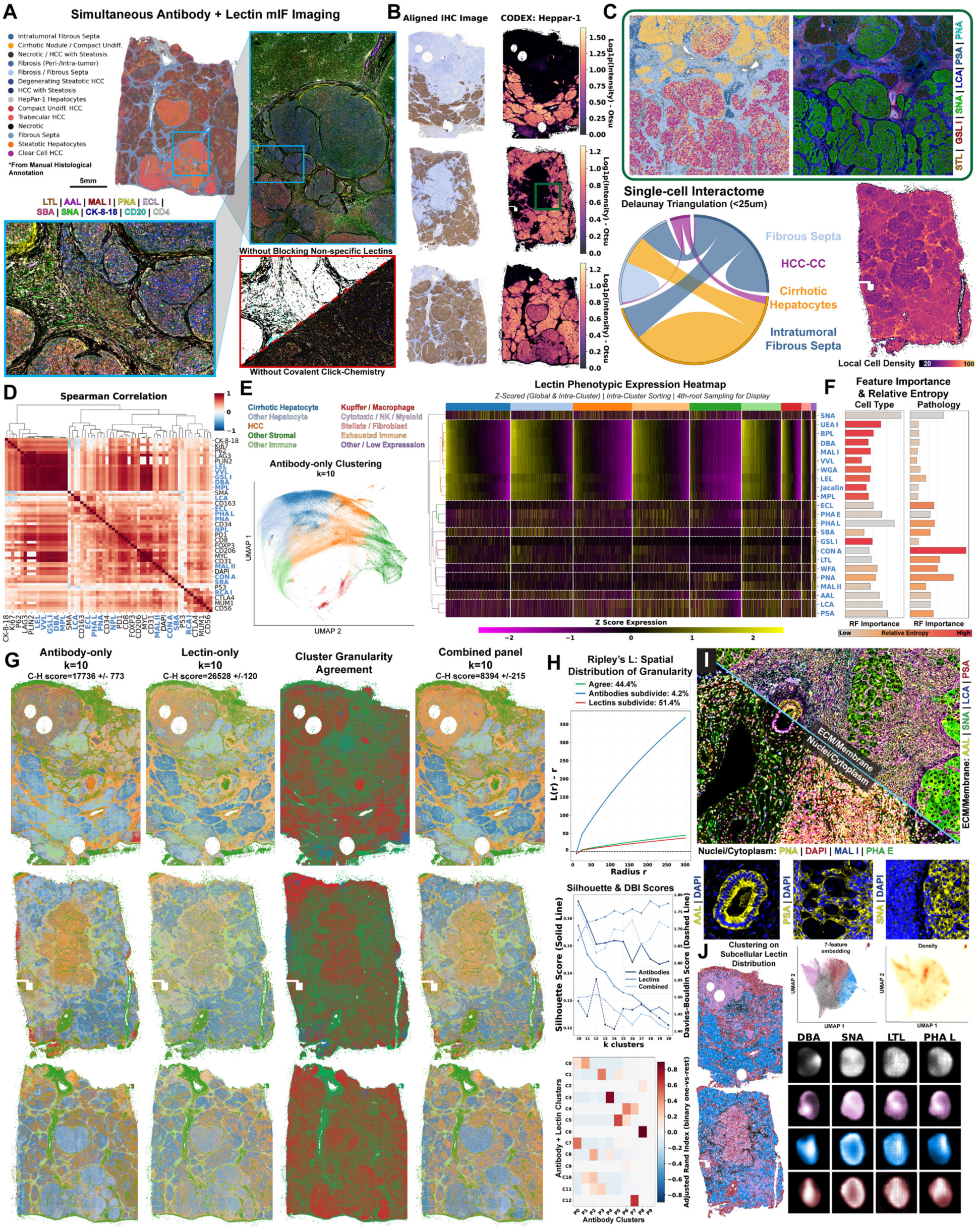
CODEX-GP for highly multiplexed imaging of glycans and proteins that reveals distinct spatial glycomic signatures at single-cell resolution. (A) Representative liver HCC biopsy with manually annotated pathology, with sequentially zoomed-in regions of interest highlight select lectin and antibody staining on the same sample. Critical considerations of blocking non-specific lectin binding and the inclusion of covalent click-chemistry barcoding are also illustrated. (B) Adjacent sections from n=3 human patients underwent IHC staining for HepPar-1, and are co-registered with the multiplex fluorescence of HepPar-1, post-lectin staining to ensure that prior lectin staining did not interfere with subsequent antibody staining. IHC HepPar-1 signal is extracted using rgb2hed. (C) Single cell validation of four major spatial compartments from figure 3, and chord diagram showing cell-type spatial interactions throughout the entire tissue. The smaller HCC population is due in part to the sparse arrangement as shown in the local cell density map. (D) Selected antibody and lectin expression correlation heatmap shows z-score spearman correlation coefficients. Some lectins and antibodies correlate more highly than others. (E) UMAP of single cell clustering on protein markers only, and manually annotated cell types. Antibody-defined cell type cluster phenotypes of selected lectins are shown, highlighting the heterogeneity of glycan signatures within each protein cluster. Cluster sizes are proportional to the 4th root of the true population sizes of each cluster. Cells within each cluster are sorted by vector-normalized average z-score intensity. (F) Random forest feature importance ranks selected lectins for both cell-type and tissue pathology classification. Bars are colored according to multi-class relative entropy. Some hierarchical families of lectins are better suited for cell type classification, while others perform better for tissue pathology classification. (G) Spatial comparison of antibody-only, lectin-only, and combination of protein and lectinK-means clustering, with cluster granularity agreement map. Green cells represent cells that are the same cluster identity in both protein-only and lectin-only clustering. Red cells represent those that are more heterogeneous in lectin-clustering than antibody-clustering. Blue cells are more heterogeneous in antibody-clustering than lectin clustering. lectin-only clustering exhibited the highest C-H score, but lowest overall Silhouette score. (H) Characteristics of the clustering differential in (G) show that cells that were more granularly subdivided by antibody markers are highly localized, while cells that are more granularly subdivided by lectins are diffuse, generally aligning with hepatocytes rather than stroma. Antibody-only clustering generally had a higher silhouette score and lower DBI score for all numbers of clusters, while lectin-only clustering had a peak silhouette score at a slightly larger number of clusters. ARI heatmap shows the subdivision of antibody clusters by lectin clusters using the optimal number of clusters defined by silhouette score. (I) Exemplary image of select lectin markers, highlighting different subcellular compartments. Selected channels highlight cell-specific labeling. (J) Subcellular distribution can also be used to cluster cell types. Spatial, UMAP and density plots are shown with select lectins and stochastic reconstructions from cVAE trained on lectin mIF images of liver tissues. Scale bar 5mm. See also Figure S13.

To validate the robustness of CODEX-GP and assess its compatibility with downstream proteins imaging, we examined HepPar-1, a classic hepatocyte marker,^35^ and found that its colorimetric DAB-based immunohistochemistry (IHC) signal in adjacent sections was consistently preserved in CODEX multiplex immunofluorescence (mIF) after lectin staining **(Figures 4B and S13)**. Furthermore, in samples MASH-HCC-1T and HCC-6T, glial fibrillary acidic protein (GFAP), a marker of hepatic stellate cells (HSCs), showed protein imaging signals specifically enriched in cirrhotic nodules without steatosis, consistent with increased hepatic stellate cell–associated signals in the cirrhotic microenvironment **(Figures S15A–S15C)**.^36^ In sample HCC-5T, CD20 protein imaging signals were specifically enriched in the fibrous septa, consistent with the marked B-cell accumulation identified by DBiT-GPT in the same region **(Figures S15A–S15C)**.

### CODEX-GP defines glycan signatures across whole tissue sections at single-cell resolution

Next, we sought to explore the glyco-codes across whole tissue sections at single-cell resolution **(Figure 4C)**. In sample HCC-5T, AAL-derived glycan signals were enriched in the NOS-HCC region, with the strongest signals observed in the right-sided HCC lesion, which exhibited a confluent multinodular architecture composed of multiple adjacent tumor nodules, whereas the left-sided early-stage HCC lesion showed weaker signals **(Figure S15A)**. The spatial distribution of AAL-binding glycans was broadly consistent with that of the co-imaged protein p62 **(Figure S15A)**. In light of previous studies showing that p62 can promote the malignancy of HCV-positive HCC through nuclear factor erythroid 2-related factor 2 (NRF2)-dependent metabolic reprogramming,^37^ this pattern suggests that AAL-recognized core fucosylation–related glycans may be associated with a more progressed tumor state in NOS-HCC. We further observed that PHA-L-derived glycans signals was highly enriched in the fibrous septa of HCC-5T **(Figure S15A)**, consistent with the DBiT-GPT sequencing results (**Figures 2F and S5B**). Previous studies have shown that PHA-L specifically binds mature MGAT5-derived glycans. MGAT5 is a golgi-resident glycosyltransferase responsible for adding GlcNAc branches to complex N-linked glycans,^38^ and PD-L1 has been identified as one of its glycoprotein substrates.^39^ In our data, PD-L1 protein were also highly and specifically enriched in the fibrous septa. Although PHA-L-binding glycans does not specifically label PD-L1, the concordant distribution of them in this region is consistent with the possibility that GlcNAc are enriched in the fibrous septal microenvironment.

PNA is among the few lectins with well-defined tumor-associated functions, as it classically recognizes the Thomsen–Friedenreich (TF) antigen, a tumor-associated glycan structure linked to malignant transformation and tumor progression in multiple carcinomas.^40^ Based on the DBiT-GPT results, PNA-binding glycans showed sample-restricted enrichment and was identified as specifically expressed only in selected samples within HCC and peritumoral fibrous stroma **(Figure 3L)**. Interestingly, CODEX-GP clearly imaged distinct spatial patterns of PNA-binding glycans in the peritumoral fibrous stroma and HCC-CC regions **(Figures S15A–S15C)**, thereby providing a systematic characterization of its spatial distribution in HCC. Similarly, SBA-binding glycans showed an NOS-HCC-associated distribution pattern in only two samples in the DBiT-GPT spatial profiling results **(Figure 3L)**. However, the imaging data demonstrated strong SBA-derived glycans signals in both cirrhotic hepatocytes and all three HCC subtypes **(Figures S15A–S15C)**, suggesting that SBA-binding glycans may reflect a broader glycan program spanning the transition from cirrhotic hepatocytes to HCC. ECL-binding glycans also did not show a clearly defined disease-associated pattern in the DBiT-GPT spatial profiling data **(Figure 3L)**. However, in the CODEX-GP imaging data, ECL-derived glycans signals were primarily enriched in the fibrous septa of HCC-5T and HCC-6T, whereas in MASH-HCC-1T they were predominantly localized to the tumor necrotic region **(Figures S15A–S15C)**.

WFA-derived glycans signals showed highly specific enrichment in the fibrous septa of sample HCC-5T **(Figure S15A)**, in agreement with the DBiT-GPT results (**Figures 2F and S5B**). However, in sample MASH-HCC-1T, WFA-derived glycans signals were mainly concentrated in the tumor region, underscoring the context-dependent and sample-specific distribution of glycan patterns across different HCC lesions. LTL-derived glycans signals was barely detectable in HCC-5T, but was highly expressed in both cirrhotic hepatocytes and HCC with steatosis in MASH-HCC-1T, as well as in cirrhotic hepatocytes and HCC-CC with steatosis in HCC-6T **(Figures S15A-S15C)**. This pattern suggests a potential association of LTL-binding glycans with steatosis-related HCC progression. Taken together, these results demonstrate that CODEX-GP imaging is broadly consistent with the glyco-codes defined by DBiT-GPT, while notably enabling more precise delineation of glycan spatial distribution, owing to its single-cell resolution across whole tissue sections.

Remarkably, both the sequencing-based and imaging-based results consistently identified PSA-binding glycans as a broad fibrosis/stroma-associated marker, with a distribution pattern more closely aligned with tumor-associated stromal regions and broadly consistent with protein alpha-smooth muscle actin (α-SMA) **(Figures 3L and S15A–S15C)**. By comparison, LCA-binding glycans showed a wider fibrosis/stroma-associated distribution, spanning fibrous septa, peritumoral fibrous stroma, and intratumoral stromal regions. These findings highlight PSA- and LCA-binding glycans as promising candidate targets for liver fibrosis and potentially for fibrotic diseases more broadly.

### Simultaneous imaging of subcellular glycans and proteins

To further systematically interrogate the co-imaged glycan–protein landscape, Spearman correlation analysis across all lectin- and antibody-derived imaging features was performed and results revealed that lectins recognizing related glycan motifs were more strongly correlated with immune, epithelial, or stromal cell markers than with other lectins **(Figure 4D)**. For instance, galactose and GalNAc binders such as SBA, PNA, and Jacalin have near-zero correlation with LCA and Con A, which are mannose binders, but correlate tightly with Collagen 1, perilipin 2 (PLIN2), CD31, and LAG3, suggesting that regions of high fibrosis and lipid accumulation exhibit elevated levels of these glycans. Despite these broad correlations, many lectin-binding glycan signatures are not necessarily cell type specific, and therefore cell type annotations of clusters can still be done manually using conventional protein markers, as shown in the UMAP, while glycans can profile cluster subdivisions within each protein-defined cluster **(Figure 4E)**. In the establishment of this new class of lectin probes for multiplex glycomics, it is critical to ascertain whether certain lectins-binding glycans are cell-type-specific, or disease-state-specific.^41^ Certain lectin-binding glycans may have more importance and statistical power in delineating protein-derived cell types, while others for pathologies, as shown in the random forest feature importance and relative entropy bar charts **(Figure 4E)**. For example, high relative entropy O-glycan or broadly distributed GlcNAc/LacNAc motif lectins such as Jacalin, UEA I, BPL, DBA, MAL I, VVL, and MPL are stably partitioned across cell types or local tissue compartments such as the epithelial, biliary, and immune cells of ductovascular anatomy, yet still show only modest random-forest importance for biopsy diagnosis since the same cell types recur across many liver diseases. In contrast, pathology-linked N-glycans implicated in fucosylation, branching, and selected exposed terminal motifs such as LTL, Con A, PNA, and PHA L may be more implicated in disease progression. Interestingly, despite LTL and UEA I being fucose binders, LTL is more tuned to terminal α1,3-fucose/Lewis x–like determinants, whereas UEA-I is more strongly associated with H-type 2/ α-fucose reactivity, potentially differentiating UEA-I for cell-type specific insight, and LTL for pathology-driven insight. These results suggest that glycans have powerful profiling capabilities that may be complementary to proteins, and as such, it is prudent to assess their individual clustering performance.

Fortunately, the simultaneous profiling of glycans and proteins preclude the need for co-registration alignment of adjacent sections and confirms that lectin-derived glycan signals are able to recover the same high-level cellular identities as proteins through unsupervised clustering **(Figure 4F)**. This concordance is stronger in some cell types than others, suggesting the role of glycome in defining cell types vs functional states. In particular, the cluster agreement map indicates that cells in stromal and normal hepatic tissue are more consistent between glycans and proteins but subdivide protein-based clusters (marked in red) particularly in tumor regions **(Figure 4G)**. This is a striking result because it is well documented that cancerous tissues exhibit metabolic dysfunction and aberrant protein dynamics. For instance, several cancers have unique subtypes, oncogenic mechanisms, and intratumoral heterogeneity as a result from proximity to vascularization, biomechanical stresses, and niche interactions with other cell types.^42–44^ Relative to protein-based clustering, glycan-based clustering exhibited a higher Calinski-Harabasz score, which measures the raw variance in the data through the ratio of the sum of between-cluster to within-cluster dispersion. However, glycan-based clustering also exhibited a lower silhouette score and higher Davies-Bouldin Index (DBI) score, suggesting that glycans yielded looser clusters since these are better measurements of geometric separation **(Figure 4G)**. Therefore, the glycans profiled in this dataset may have a larger variance than the proteins included, and potentially stratify cells along a different, potentially functional metabolic, dimension rather than a highly specific cell-type discretizing one **(Figure 4H)**.

In addition to cell-average clustering, lectin-binding glycans also exhibit unique subcellular localizations, similar to many proteins. In the validation of this new class of multiplexing probes, target characterization and validation may be of significant interest, especially for future flow-cytometric cell sorting using these probes. For instance, a membrane marker may not be as useful for single nucleus sequencing preparation. **Figure 4I** illustrates select glycan markers and their nuclear or perinuclear localizations versus membrane and extracellular localizations. Importantly, different functional tissue units such as the bile ducts, stromal septa, and hepatic nodules shown, have different glycans that localize to the cytoplasm, indicating some level of cell-type specificity. Leveraging this subcellular spatial dimension, we can even cluster cell types into broad structures resembling expected anatomy. Training a cVAE on 10 million cells reveals archetypal subcellular distributions of each broad cluster, reconstructed from stochastic representations of the subcellular spatial distribution embedding space **(Figure 4J)**. These initial tests confirmed the performance and efficacy of simultaneous glycan and protein mIF and revealed critical characterizations of lectin-derived glycan imaging such as their complementary information and subcellular distribution utility. The rich single-cell and even subcellular spatial information afforded by this technology can add color to the multi-omics landscape and be useful in previously unexplored ways for translational impact across many diseases.

### CODEX-GP reveals glyco-codes of liver disease

We further applied high-throughput CODEX-GP imaging to clinical liver tissue cores from 302 patients with various diagnoses **(Figure 5A)**. Representative raw CODEX images of selected diseases are shown with select lectin-binding glycan markers **(Figure 5B)**. Mean image intensity values derived from antibody and lectin signals for all cells were pooled and clustered using antibody signals exclusively for protein-based cell-type annotations, and lectin signals exclusively for glycan signature clusters, or glyco-codes. The composition of these glyco-codes at the cellular and tissue levels are shown in the Sankey diagram **(Figure 5C)**. All marker intensities were used to generate UMAPs colored according to the cell-type annotation **(Figure 5D)** and pathologist clinical classification of the tissue microarray (TMA) core from which those cells were derived **(Figure 5E)**. To understand how these glyco-codes interface with protein markers, the glyco-code and protein cluster for each cell is mapped using Delaunay triangulation. The log_2_-odds for the interaction of a given protein cluster and glyco-code are mapped in a network diagram **(Figure 5F)** and show how some clusters are within 1 physical degree of freedom many orders of magnitude more frequently than by random chance. Three such cluster neighborhood interactions stood out; an “Immune Evasion”, a “T-Cell Exhaustion”, and a “Steato-Fibrotic” niche.

**Figure 5.**
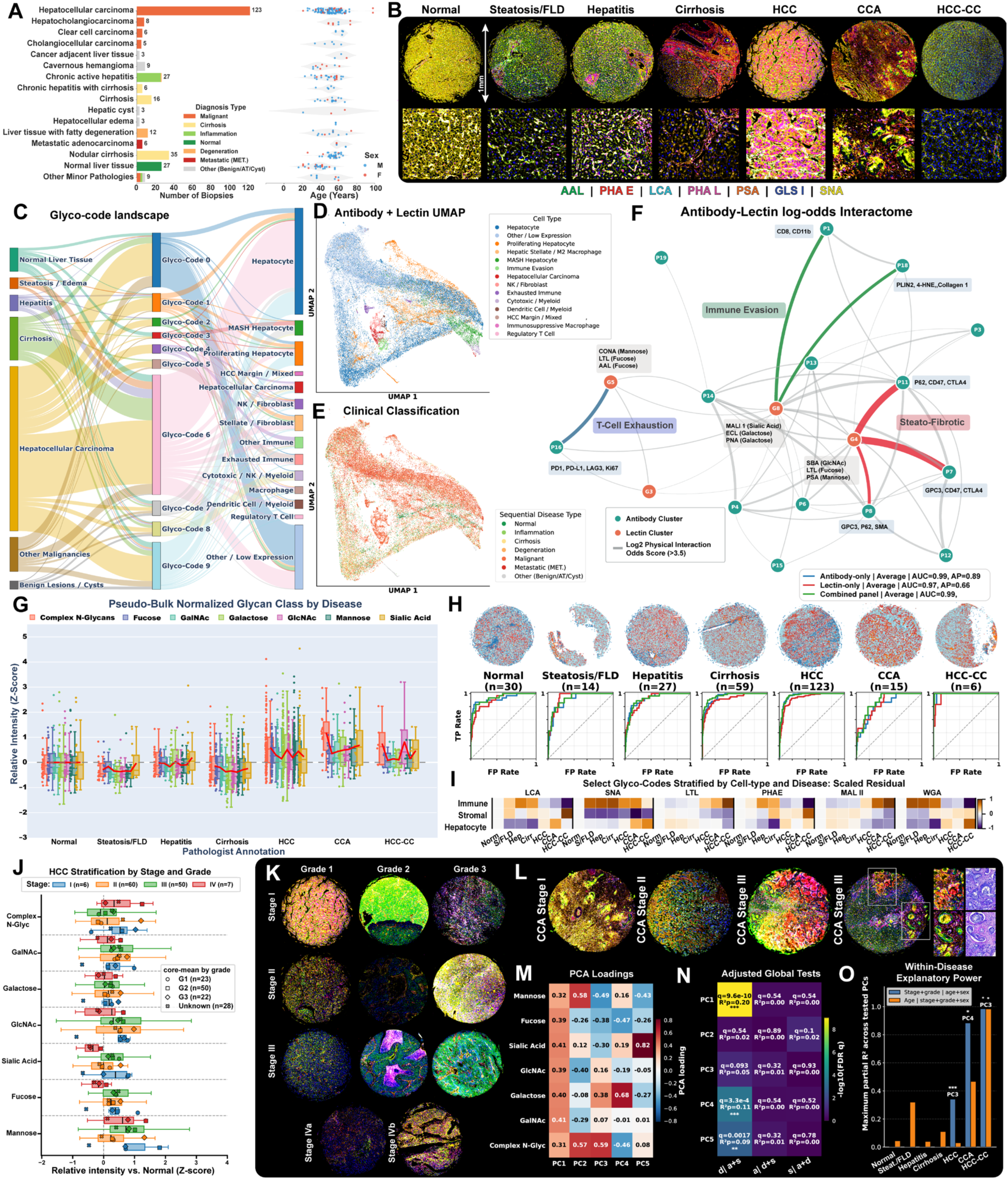
CODEX-GP imaging of liver disease tissue cores from 302 patients revealed disease-specific glyco-codes as potential biomarkers and key molecular mechanisms. (A) Summary of liver disease tissue core dataset. (B) Raw CODEX images of select lectins in disease-specific exemplary cores. (C) Separate antibody-only and lectin-only clustering Sankey diagram shows the composition of manually annotated cell types and tissue types for each lectin cluster. (D-E) UMAPs of cells from the TMA dataset clustered by antibodies + lectins combined markers colored according to protein-only cell-type (top) and clinical annotation of the cores from which those cells were derived (bottom). (F) Log2 odds interactome between antibody (i=20) and lectin (j=10) clusters. Only cluster interactions enriched beyond 2^3.5 are shown. Edge weight is proportional to log2 odds. Selected interactions are highlighted and annotated. (G) Box and swarm plots of z-scored core mean lectin expression for certain classes of lectins. Class-level means are shown in red lines. (H) One-versus-rest classification performance summary of several liver diseases show the average AUROC and AP of 7 classification methods. (I) Normalized z-scored residuals of select lectins for each pathology, stratified by key protein-defined cell-types highlight the importance of simultaneous protein and lectin profiling. (J) Stratification of HCC cores by pathologist annotated stage, with exemplary same-slide CODEX images of different stages of HCC, as manually annotated by pathologists. (K) Raw CODEX images of the lectins previously described in (B) highlighting heterogeneity among disease stages and grades. (L) Raw CODEX examples of hepatocholangiocarcinoma (CCA) differences across stages, with an exemplary core of stage 3 CCA exhibiting inflamed vasculature easily detectable using lectins and less apparent through histological staining. (M) Loading heatmap for the first five PCs illustrates the relative contribution of each glycan class to the primary axes of variation. (N) Covariate-adjusted global regression analysis assesses the explanatory power of disease group, age, and sex for each PC. Heatmap cells report Benjamini–Hochberg FDR-adjusted q-values and partial coefficients of determination (R²p) derived from nested ordinary least-squares regression models. Models were tested by sequentially adding disease group, age, or sex to evaluate their independent contributions. (O) Within-disease regression summary compares the maximum partial R² for stage/grade effects against age effects across all tested PCs. Bars represent the largest partial R² observed for each group, with asterisks denoting FDR-adjusted significance for the specific PC. Statistical significance was determined using partial F-tests and controlled via the Benjamini–Hochberg procedure. Significance thresholds: * q < 0.05, ** q < 0.01, and *** q < 0.001. See also Figure S14.

The “Immune Evasion” niche had the strongest interactions in the Dataset (LogOdds: 6.4 to 10.3). These clusters express relatively high levels of GPC3 (the hallmark HCC tumor marker), CD47 (the “don’t eat me” macrophage evasion signal), CTLA4, and p62 (autophagy/stress). The glycans in this neighborhood are defined by intense expression of SBA (GalNAc), LTL (Fucose), and PSA (Mannose). These clusters are mainly localized in HCC tumor beds (as defined in **Figures 3D and 4C**). In clinical biology, hyper-fucosylation (marked here by LTL) is a well-known driver of HCC (e.g., the FDA-approved AFP-L3 test measures fucosylated AFP). Results suggest these Fucose / GalNAc-rich glycan structures may act as ECM anchors for the tumor cells that are actively deploying CD47 and CTLA4 to shut down local Kupffer cells and immune surveillance.

The “T-Cell Exhaustion” niche (LogOdds: 6.8) highly expresses the suppressive immune checkpoints PD-1, PD-L1, and LAG3, alongside Ki67 (proliferation). These cells are also dominated by Con A (Mannose), LTL (Fucose), and AAL (Fucose). Exhausted T-cells and suppressive myeloid cells may not be randomly distributed but are found concomitantly in heavily fucosylated and mannosylated niches. Macrophages and dendritic cells use specific lectin receptors (like MR or DC-SIGN) to bind to mannose and fucose, which often triggers immunosuppressive cytokine release. The highly expressed glycans in these cells may act as a physical “off-switch” sponge, capturing and exhausting PD-1+ immune cells before they can reach the tumor.

The “Steato-Fibrotic / MASH” niche (LogOdds: 6.3 to 8.4) express some hallmarks of MASH such as PLIN2 (lipid droplets/steatosis), 4-HNE (lipid peroxidation/oxidative stress), and Collagen I (active fibrosis). Additionally, there is moderately high expression of cytotoxic/myeloid infiltrates (CD8, CD11b). This cluster is also dominated by MALII (α-2,3 Sialic Acid), ECL (Galactose), and PNA (Galactose / T-antigen). As the liver transitions from fatty liver to cirrhosis, HSCs lay down dense collagen tracks. The physical co-localization of Collagen I / 4-HNE with Sialic Acid (MAL II) and exposed Galactose (ECL/PNA) in this dataset suggests that the de-sialylation of the tissue (exposing underlying galactose residues) is a structural hallmark of the fibrotic MASH state. It is along these specific sialic/galactose boundaries that CD8+ and CD11b+ immune infiltrates may get bottlenecked.

Naturally, due to the relatively predictable nature of liver cancer progression, it is critical to ascertain the glyco-codes of various disease stages to establish lectin-based multiplex imaging as a viable method for early detection and furthering our understanding of these disease mechanisms. Lectin markers were grouped in broad glycan classes for clarity, and mean intensities for each core are normalized and plotted **(Figure 5G)**. The mean intensity of these glycan classes is outlined in red for each disease category, but there is significant variability within each disease. While on average, one might expect broad shifts in lectin-derived glycan imaging intensity to correlate with certain diseases such as an overall diminished intensity for steatosis and cirrhosis, the actual classification utility of lectins may suffer at this scale. To avoid leakage from spatially autocorrelated neighboring cells within the same TMA core, classification was performed on core-level aggregated features rather than individual-cell measurements, and all cells from a given core were kept within the same cross-validation fold. Using core-level feature vectors across seven classification architectures (**Figure S14**), the average AUROC and average precision (AP) indicated that, for most disease states except hepatitis, glycans alone are sufficient to distinguish various pathologies despite slightly lower performance as compared to multiplex protein imaging. This is not surprising since the protein markers were specifically chosen for detecting these liver disease states while the 23-plex lectin panel may serve as a universal multiplex biomarker platform for pan-liver disease diagnostics. Interestingly, for Hepatocholangiocarcinoma (CCA), the combination of glycans and proteins performed even best. Nevertheless, the lectin-based glyco-codes proved to be decent biomarkers for these diseases, achieving AUROCs within 10% of the antibody-based protein gold standards across all disease states. When thresholding for specific cell types using protein markers within each disease category, immune, stromal, and hepatocyte cells exhibited unique lectin-based glycan profiles **(Figure 5I)**. Together, this suggests yet again that glycans may be stratifying along a dimension orthogonal to proteins, and classification performance may suffer from biased diagnoses that were based on IHC staining with antibodies. Therefore, we stratified the data from **(Figure 5G)**, specifically the HCC TMA cores, which showed the largest standard deviations in glycan class intensities, along pathologist annotated stage and grade **(Figure 5J)**. Exemplary raw CODEX images using the same lectin markers as in **(Figure 5B)** highlight the heterogeneity between grade, stage, and region of interest **(Figure 5K)**. This heterogeneity may be the source of this wide standard deviation even within each disease category and suggest that lectins may be suitable for high-content granular delineation of disease progression and early detection. To highlight this, CCA cores of different stages are shown **(Figure 5L)**, where the last stage III core exhibits inflamed vasculature consistent with both stage I and stage III phenotypes that are less apparent in the routine histology acquired on the same sample post CODEX. However, the caveat is that age could be a confounding factor, as shifts in overall lectin intensity may correlate with age in certain disease groups. This is significant because age and disease stage are not wholly independent, as more advanced stages of disease manifest in older individuals on average. This co-linearity may therefore diminish the utility of glycans as biomarkers for stage-level granularity. To test this, mean glycan core intensities were pooled and converted to principal components (PC). In the 7 disease categories, the top 5 PCs encompassed 85% of the total variance, and their glycan class loadings are shown in **(Figure 5M)**. Multiple regression testing of pooled glycan class data shows most disease-specific glyco-code changes are attributable to broad global changes (PC1) and are largely independent of age or sex **(Figure 5N)**. For HCC especially, disease grade and stage had significantly higher explanatory power for the source of variance **(Figure 5O)**. Together these data suggest that lectin-based highly multiplex glycomic imaging can indeed be a powerful tool to profile liver disease and may unlock an easily distinguishable and interpretable dimension previously underexplored. A closer look at “normal” TMA cores that were misclassified using lectin markers reveals that despite being relatively indistinguishable through histological staining or antibody markers, they had a much stronger classification confidence for hepatitis, driven by canonical glycan signatures for inflammation such as LTL, LEL, and SBA (**Figure S14**).

## DISCUSSION

We developed spatial-GPT as a spatial tri-omics platform for the simultaneous profiling of glycans, proteins, and/or transcripts. Although recent advances have substantially improved RNA capture from FFPE tissues,^45,46^ severe RNA degradation remained a limitation in clinical archival tissues stored for a long period of time and, in some samples, prevented robust recovery of informative spatial transcriptomic features. In contrast to RNA, glycan features appeared markedly more stable in archived FFPE tissues, likely because fixation suppresses endogenous enzymatic activity,^47^ while lectin-based detection can still capture preserved glycan motifs even when partial degradation has occurred.^41^ As a result, glycan mapping remained highly robust even in archived FFPE tissue blocks stored for over 8 years and consistently recapitulated histopathological architecture. This feature substantially expands the biological information that can be extracted from clinical FFPE specimens, particularly from ultra-long storage time archival blocks. The CODEX-based co-imaging strategy for glycans and proteins further provides a high-throughput platform for full-section spatial mapping at single-cell and subcellular resolution, whereas DBiT-GPT adds the total RNA whole-transcriptome layer that is missing from CODEX-GP and is essential for mechanistic exploration. At the same time, because of its distinct chemical labeling strategy and broader compatibility, DBiT-GPT can be more readily adapted for further modification and capture of diverse glycan motifs. Together, these two platforms are highly complementary and synergistic.

In this study, a lectin-based approach enabled highly multiplexed spatial mapping of functionally relevant glycan motifs rather than whole glycan structures in situ. Such motif-level information is particularly valuable because it is more directly implicated in functions and actionable for downstream applications, enabling strategies such as blocking glycan-mediated signaling via lectin binding,^48–50^ targeted glycan trimming,^51^ or modulation of glycosyltransferases involved in generating the corresponding glycan motifs, for example MGAT5 for PHA-L–reactive branched N-glycans, FUT8 for core fucosylation, ST6GAL1 for α2,6-sialylation, and B4GALT1 for RCA I–reactive terminal galactose motifs.^52–55^ To date, lectin-based spatial glycan technologies have been limited to direct imaging of lectin probes, whereas DBiT-GPT is the first platform to enable glycan profiling via spatial barcode sequencing. Meanwhile, existing lectin-based imaging approaches are typically limited to a handful of lectins,^56,57^ and only very recent studies have extended this number to six.^58^ By contrast, CODEX-GP incorporates an innovative design and workflow that simultaneously maps the binding of 23 lectins, with the potential to be expanded to an even larger panel. Therefore, DBiT-GPT and CODEX-GP provide a uniquely complementary framework for systematic spatial glycan capture at both the sequencing and imaging levels.

Spatial-GPT was developed as a transformative and integrative platform for interrogating tissue biology. By co-profiling molecular layers across an extended central dogma, spatial-GPT enabled not only the identification of differentially expressed glycans, proteins, and transcripts across distinct stages of HCC progression, but also the discovery of biologically meaningful crosstalk among them. For example, in both NOS-HCC and HCC with steatosis, we identified glycans together with glycosyltransferase genes predicted to generate the corresponding glycan motifs and found concordant spatial distributions and expression trends between them. In vitro validation further showed that knockdown of glycosyltransferase genes associated with broadly enriched tumor glycans reduced tumor colony formation. These results suggest that targeting glycosyltransferases may offer translational potential as compared to direct lectin blockade, particularly because lectins may exhibit moderate to strong immunogenicity.^59^ Moreover, spatial-GPT also enabled glycan–protein colocalization analysis. In DBiT-GPT, we identified strong colocalization of GSL I with CXCR3 and CX3CR1 in intratumoral fibrous septa regions, whereas CODEX-GP revealed colocalization of PHA-L with the immune checkpoint protein PD-L1. Recent studies support the therapeutic potential of combining lectin-based glycan targeting with immune-directed modalities. Antibody–lectin chimeras (AbLecs) and glycan-dependent T cell recruiters (GlyTRs) have shown that lectin-mediated recognition of tumor-associated glycans can be coupled with antibody or T cell–engaging functions to enhance anti-tumor activity.^60,61^ Consistent with this concept, inhibition of FUT8-mediated core fucosylation has been shown to reduce cell-surface PD-1 expression and enhance T cell activation.^62^ Together, these findings suggest that once robust glycosylation patterns on immune checkpoints are defined, glycan-targeted modulation may be combined with checkpoint blockade to elicit stronger anti-tumor immune responses.

Finally, high throughput imaging-based spatial glycan-protein co-profiling across more than 300 liver disease tissue specimens enabled the systematic identification of glyco-codes associated with distinct stages of HCC progression and different HCC subtypes. These results provide a foundation to develop single-cell or spatial glyco-codes as potential biomarkers and nominating novel targets for glycotherapeutics.

### Limitations of the study

In this study, the glycan-detection panel did not include Siglecs (Sialic acid-binding immunoglobulin-type lectins), a subset of I-type lectins that are primarily expressed on immune-cell surfaces and function as well-characterized sialic acid-binding immunoglobulin-like co-receptors.^63^ Instead, our 30-lectin panel was designed as a systems-level spatial glycomics platform to map diverse glycan classes and uncover previously unrecognized glyco-codes associated with tumor progression and heterogeneity. Future studies focused on glycan-mediated immune regulation could incorporate Siglecs and other mammalian glycan-binding lectins to directly assess receptor-specific glycan interactions in the tumor microenvironment. The DBiT-GPT spatial sequencing data generated in this study are not at near-single-cell resolution. Because human liver specimens are relatively large and contain spatially separated regions representing different stages of disease progression, we used a chip with a larger capture area to maximize tissue coverage. This design necessarily reduced spatial resolution but enabled simultaneous profiling of multiple disease-associated tissue regions within the same section. In future studies, this limitation could be addressed by applying methods such as SuperFocus,^64^ a modality-agnostic computational platform that integrates histopathology with pixel-based spatial measurements to generate single-cell spatial multi-omics maps. In parallel, we partially mitigated this limitation by applying CODEX-GP to adjacent tissue sections, which provided subcellular-resolution glycan–protein co-imaging and enabled cell- and tissue-architecture-level validation of key spatial patterns identified by DBiT-GPT.

## RESOURCE AVAILABILITY

### Lead contact

Further information and requests for resources and reagents may be directed to the lead contact, Dr. Rong Fan.

### Materials availability

All materials used for Spatial-GPT are commercially available.

### Data and code availability

Raw and processed sequencing data generated in this study were deposited in GEO under accession number GSE333851. CODEX imaging datasets are available at Zenodo: HCC-5T, 10.5281/zenodo.20130510; HCC-6T, 10.5281/zenodo.20128058; CODEX_TMA_LV1201b, 10.5281/zenodo.20140699; CODEX_TMA_BC03117a, 10.5281/zenodo.20144328; and CODEX_TMA_LV961, 10.5281/zenodo.20144918. The raw imaging data for MASH-HCC-1T exceed 50 GB and are therefore too large to be deposited in the public repository; these data will be stored on a private cloud-based server and made available by the corresponding authors upon reasonable request.

All data were analyzed with standard programs and packages, as detailed above. Custom code supporting this study is available at https://github.com/XiaolongTianLab/Spatial-GPT. Additional information required to reanalyze the data reported in this paper is available from the lead contact upon request.

## ACKNOWLEDGMENTS

We thank Dr. Jun Lu at Yale University for helpful discussions. We acknowledge Yale Pathology Tissue Services (YPTS) for assistance with sample preparation and histological, histochemical, and immunohistochemical staining. We thank the Yale Center for Genome Analysis (YCGA) for providing computational resources and Novogene USA for sequencing support. The molds for microfluidic devices were fabricated at the Yale University School of Engineering and Applied Science (SEAS) Nanofabrication Center. This research was supported by the US National Institutes of Health (U54CA274509, U54CA268083, UH3CA257393, RF1MH128876, U54AG079759, U54AG076043, R01CA245313, RM1MH132648 to R.F. and U01CA294514 to R.F., M.L.X. and Z.M., and the Boehringer Ingelheim Pharma (GR130134).

## AUTHOR CONTRIBUTIONS

X.T., R.F., and X.S. conceived the project, and R.F. supervised the study.

X.T. developed the technology and led the experimental work with assistance from X.S., B.C., M.Z., B.T., and X.L.

X.T., A.F., and D.Z. performed data analysis with assistance from K.L., Y.D., Y.L., X.S., M.Y., A.B., A.I., and F.G.

C.L. and X.Z. provided HCC samples and contributed to pathological assessment and interpretation.

L.Z. and B.C. conducted the in vitro validation experiments.

X.T. and A.F. drafted the manuscript with critical input and editing from R.F.

R.F., C.L., X.Y., M.G., M.X., F.L., and S.C. provided scientific guidance and reviewed the manuscript. All authors reviewed and approved the final manuscript.

## SUPPLEMENTAL INFORMATION

**Document S1. Figures S1–S15 and Tables S1**

**Table S2. DBiT-GPT reagents and oligos.**

**Table S3. CODEX-GP reagents and oligos.**

## STAR★METHODS

## KEY RESOURCES TABLE

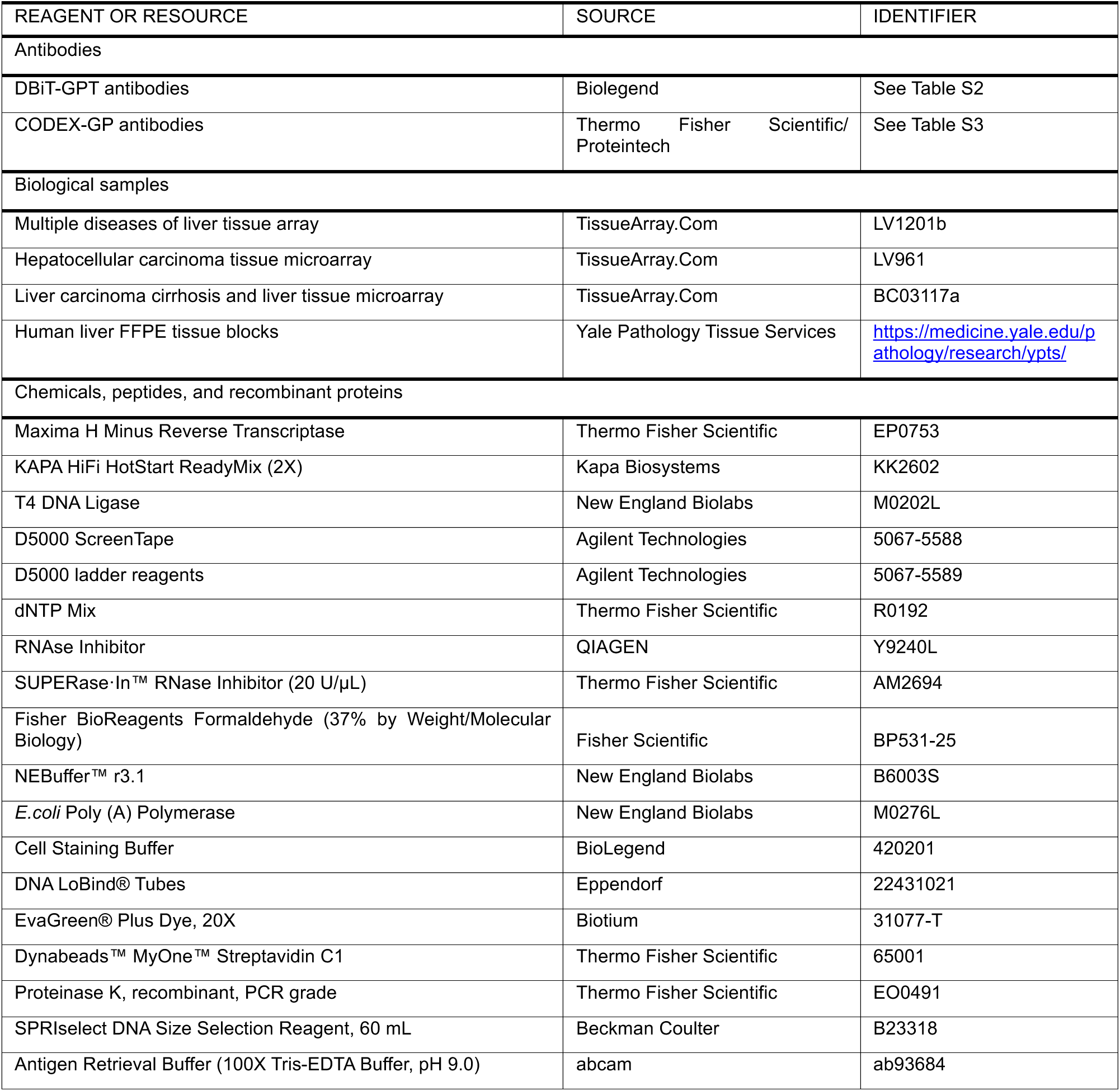
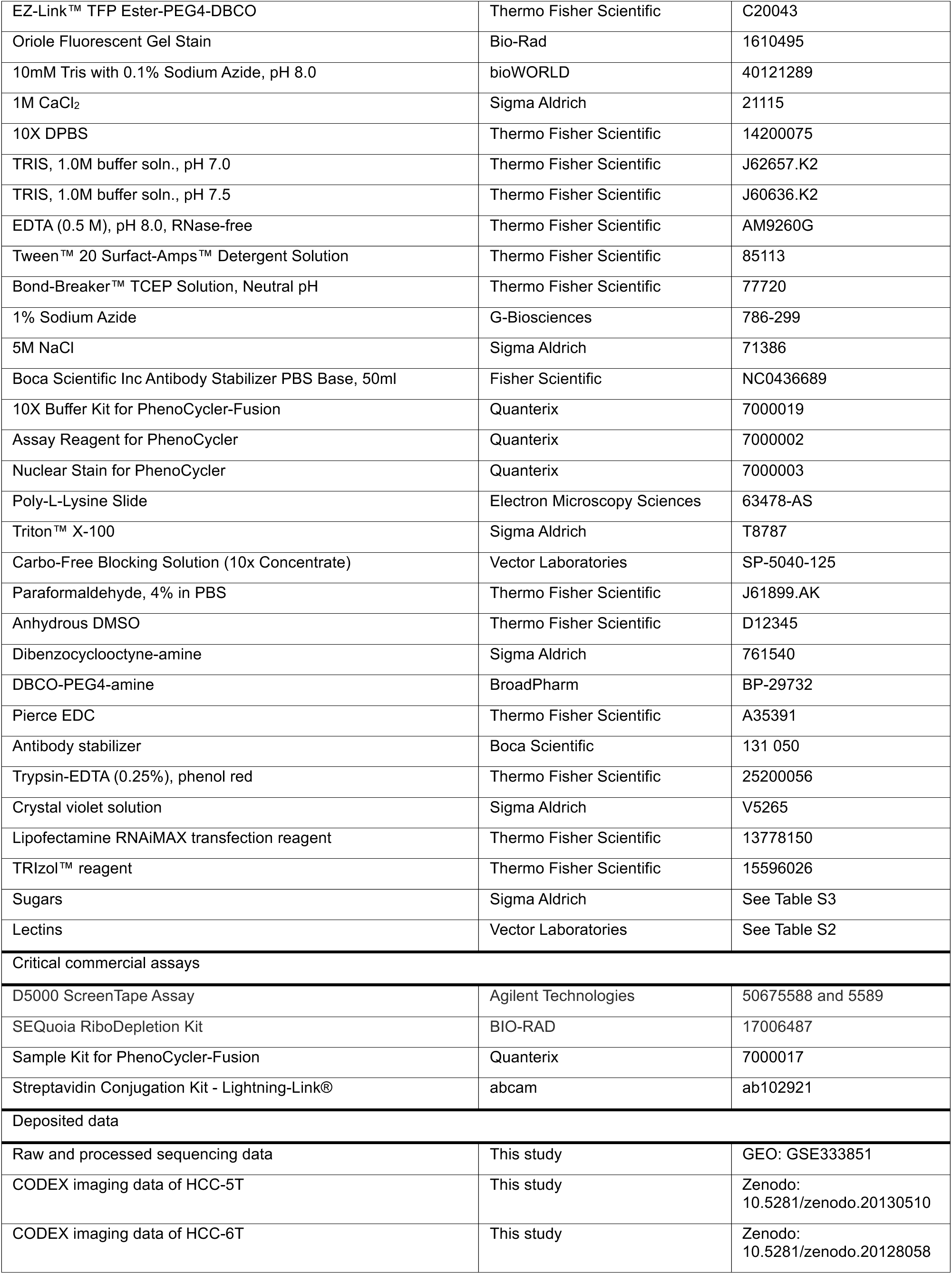
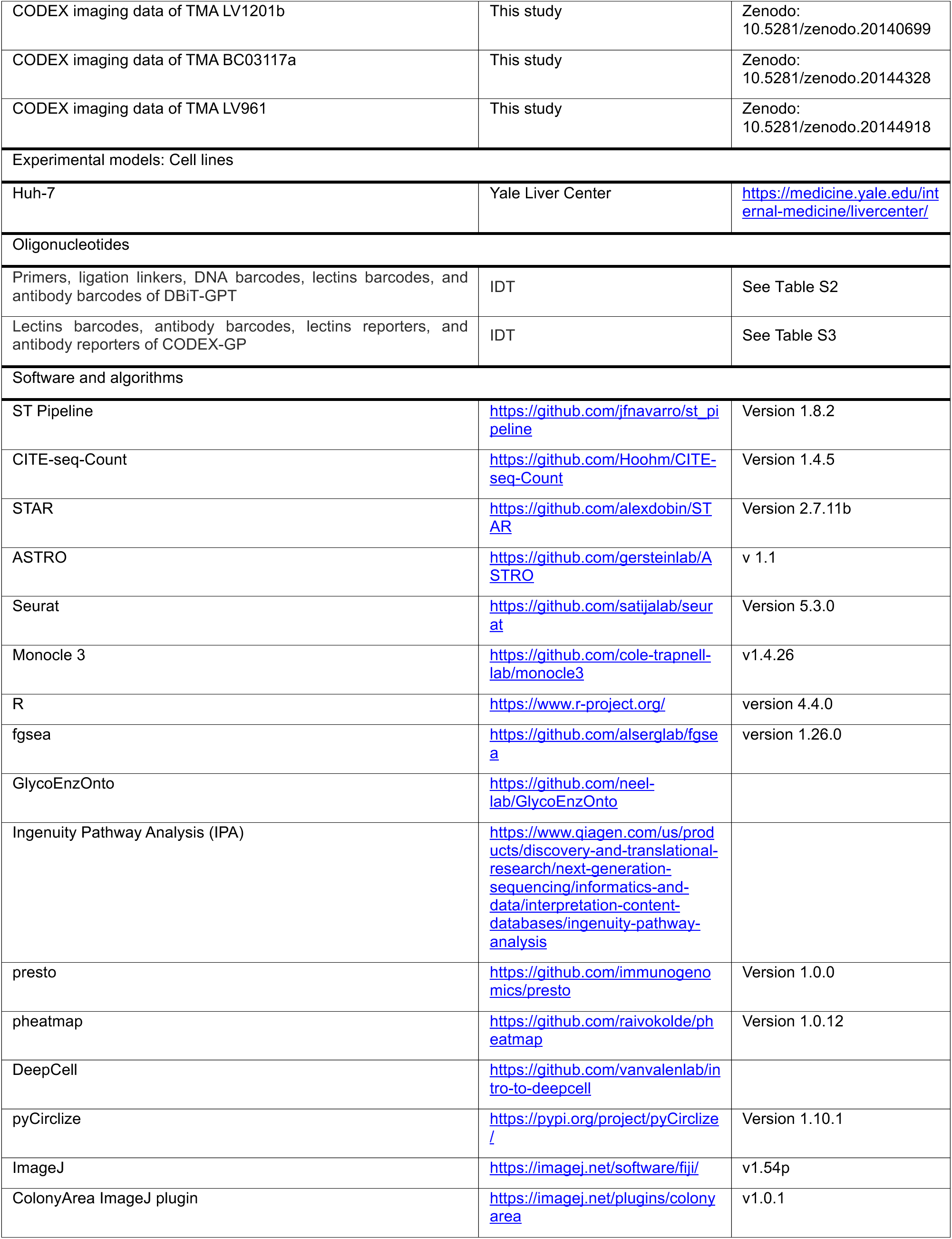
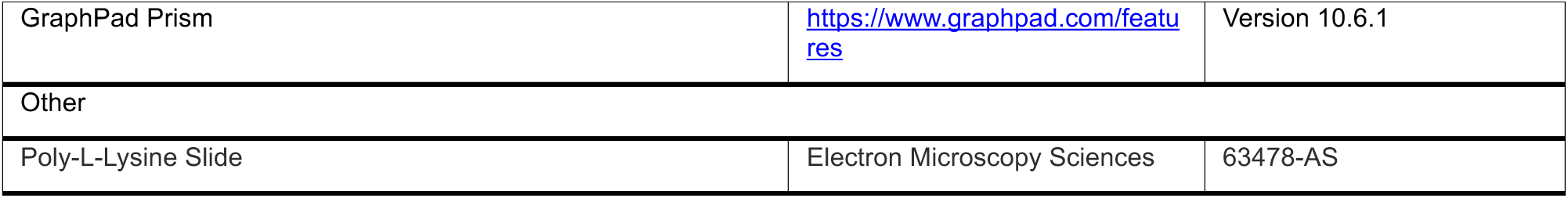

### Human HCC specimens

De-identified archival FFPE human liver tissue blocks were obtained from YPTS. The samples included 6 conventional HCC tumor tissues (HCC-T), 4 background non-tumor liver tissues from the same HCC patients (HCC-N), and 6 metabolic dysfunction-associated steatohepatitis-derived HCC tissues (MASH-HCC) **(Figure S1A)**. These tissues had originally been collected by physicians for diagnostic purposes. All tissue collection and handling procedures were conducted under approval of the Yale University Institutional Review Board, with oversight from the Tissue Resource Oversight Committee. All specimens were managed in compliance with HIPAA regulations, University Research Policies, Pathology Department diagnostic requirements, and Hospital by-laws. FFPE tissue blocks were sectioned at a thickness of 5-10 μm and mounted at the center of Poly-L-Lysine-coated glass slides (Electron Microscopy Sciences, 63478-AS). Serial sections were collected for DBiT-GPT, CODEX-GP, and additional staining assays. Sectioning of the HCC patient samples was performed by YPTS team, and the paraffin sections were immediately stored at -80 °C until further use.

### TMAs

Multiple diseases of liver TMAs (LV1201b, LV961, and BC03117a) were purchased from TissueArray.Com. These TMAs contained cores representing a spectrum of liver pathologies, including normal liver tissue, fatty degeneration, hepatitis, liver cirrhosis, cyst, cavernous hemangioma, cancer-adjacent liver tissue, hepatocellular carcinoma, cholangiocellular carcinoma, and mixed carcinoma. All TMA cores were FFPE samples sectioned at a thickness of 5 μm, and the TMA slides were stored at -80 °C until further use.

### Cells

The human hepatocellular carcinoma cell line Huh-7 was obtained from the Yale Liver Center. Cells were cultured in DMEM supplemented with 10% FBS (Invitrogen) and maintained at 37°C in a humidified incubator containing 5% CO_2_.

### Reagents and oligonucleotides (oligos)

All commonly used reagents in this study are summarized in **Table S1**. **Table S2** lists the reagents and oligos used for DBiT-GPT spatial multi-omics sequencing, including lectins, antibodies, primers, spatial barcodes, lectin barcodes, and antibody barcodes. **Table S3** lists the reagents and oligos used for CODEX-based spatial imaging of glycans and proteins, including inhibitory sugars, antibodies, lectin barcodes and reporters, and antibody barcodes and reporters.

### H&E, IHC, PAS, and diastase–periodic acid–Schiff (D-PAS) staining

H&E, IHC, PAS, and D-PAS staining in this study were performed at YPTS. All procedures were conducted in accordance with Clinical Laboratory Improvement Amendments (CLIA)-certified laboratory protocols and standard YPTS procedures.

### Preparation of DNA-barcoded lectins for DBiT-GPT

Biotinylated DNA barcodes **(Table S2)** for DBiT-GPT were conjugated to lectins using the Streptavidin Conjugation Kit - Lightning-Link® (abcam, ab102921). Briefly, for the kit with a 100-µg product size, 0.67 nmol of each unconjugated lectin was diluted in 100 µL of lectin dilution buffer (20 mM HEPES, 150 mM NaCl, and 0.1 mM CaCl₂, pH 8.5) according to the manufacturer’s instructions. Next, 10 µL of modifier reagent was added to the unconjugated lectin solution and mixed gently. The modified sample was then transferred directly into the vial containing the lyophilized streptavidin conjugation mix, gently pipetted up and down once or twice to fully resuspend the material, and incubated for 3 h at room temperature in the dark. After conjugation, 10 µL of quencher reagent was added to the lectin solution, mixed gently, and incubated for 30 min before use.

### DBiT-GPT

FFPE tissue sections at 10 μm thickness were retrieved from -80 °C freezer and equilibrated to room temperature for 10 min. Once the surface moisture had fully dissipated, the slides were then baked at 60 °C for 1 h to soften the paraffin and enhance tissue adhesion. Deparaffinization was carried out by immersing the slides in xylene twice, followed by sequential rehydration in 100%, 90%, 70%, and 50% ethanol and a final rinse in distilled water. Each incubation step was performed for 5 min. The slides were immediately placed in staining jars containing preheated 1× antigen retrieval buffer (abcam, ab93684), steamed for 30 min, and then cooled on ice for an additional 30 min to room temperature.

For permeabilization, the tissue section was covered with 500 μL of 0.5% Triton X-100 (Sigma Aldrich, T8787) in PBS and incubated at room temperature for 20 min. The section was then washed with 0.5× PBS-RI (0.5× PBS with 0.05U/μL RNase Inhibitor in nuclease-free water).

In situ polyadenylation was then performed. Briefly, a polydimethylsiloxane (PDMS) reservoir was tightly mounted onto the slide, and 60 μL of Poly(A) enzymatic mix, prepared according to the Patho-DBiT protocol,^10^ was added to the reaction chamber. The slide was incubated at 37 °C for 25 min, followed by three washes with 0.5× PBS-RI.

For lectin staining, the tissue was first covered with 100 μL of 1× Carbo-free blocking solution (SP-5040-125, Vector Laboratories) and incubated at room temperature for 30 minutes. After three washes with PBS, 50 μL of DNA oligo-conjugated lectin cocktail was applied to the tissue and incubated at room temperature for 1 hour. Unbound lectins were removed by three times wash with 0.5× PBS-RI. To block excess binding sites on tissue-bound lectins, sections were fixed with 1% Paraformaldehyde (PFA, Thermo Fisher Scientific, J61899.AK) for 5 min at room temperature, followed by incubation with 100 μL of sugar mixture (Table S3) at 4°C for 10 min. The sugar mixture was then replaced with fresh solution, and sections were incubated for an additional 10 min at 4°C according to the manufacturer’s instructions. Sugar concentrations are listed in Table S3. Tissue sections were then washed three times with 0.5× PBS-RI.

Next, the tissue section was blocked with 100μL cell staining buffer (BioLegend, 420201) with 0.05U/μL RNase inhibitor (QIAGEN, Y9240L) at 4 °C for 10 min. After three washes with 0.5× PBS-RI, the ADT cocktail was added to the tissue and incubated at 4 °C for 30 min. Unbound antibody cocktail was then removed by three times wash with 0.5× PBS-RI.

Next, the reverse transcription (RT) mixture for each tissue section was prepared in a total volume of 60 μL, containing 12 μL of 5× Maxima RT buffer, 15 μL of 30% PEG 8000, 22 μL of 100 μM 15T biotinylated RT primer, 6 μL of Maxima H Minus Reverse Transcriptase (200 U/μL), 4 μL of dNTPs (10 mM), 0.6 μL of SUPERase•In RNase Inhibitor (20 U/μL), and 0.4 μL of RNase inhibitor (40 U/μL). The RT mixture was loaded onto the tissue section using a PDMS reservoir, followed by incubation at room temperature for 30 min and then at 42°C for 90 min. After three times wash with 0.5× PBS-RI and rinse in nuclease-free water, the tissue section was imaged using a 10× objective on EVOS M7000 Imaging System (Thermo Fisher Scientific, Waltham, MA, USA).

The subsequent steps, including spatial barcode A and barcode B (Table S3) ligation, tissue section lysis, cDNA extraction, template switching, and library preparation, were performed as previously described in the Spatial-CITE-seq protocol.^11^

### Preparation of DNA-barcoded lectins for CODEX-GP

Given that streptavidin-biotin conjugation strategy used for DNA-barcoded lectins in DBiT-GPT is non-covalent, its stability may be reduced during repeated dimethyl sulfoxide (DMSO) wash steps across CODEX cycles. Therefore, a click chemistry-based strategy^34^ that generates a covalent linkage was used for preparation of DNA-barcoded lectins for CODEX-GP. In brief, EZ-Link TFP Ester-PEG4-DBCO (TFP-DBCO, Thermo Fisher Scientific, C20043) was first equilibrated to room temperature for at least 15 minutes before use. All lectins **(Table S3)** were first diluted to the same molar concentration in lectin dilution buffer. TFP-DBCO was then dissolved in anhydrous DMSO (Thermo Fisher Scientific, D12345) and added to the diluted lectins at a 1:10 lectin-to-DBCO molar ratio. The reaction mixture was incubated at room temperature for 2 hours in the dark.

For lectins showing low TFP-DBCO conjugation efficiency, an alternative EDC-based strategy was used. Briefly, lectins to conjugate were first buffer exchanged into freshly prepared EDC conjugation buffer (100 mM MES, 150 mM NaCl, pH 5.0) using a 10 kDa centrifugal filter (Sigma Aldrich, UFC501096) through three rounds of washing. DBCO-PEG4-amine (BroadPharm, BP-29732) was first dissolved in anhydrous DMSO to 100 mM and was then added for each buffer-exchanged lectin to reach a 250-fold molar excess relative to lectin. EDC solid (Thermo Fisher Scientific, A35391) was equilibrated to room temperature in the dark for at least 15 min before opening and then dissolved in EDC conjugation buffer. After complete dissolution, EDC solution was immediately added to the lectin/DBCO-PEG4-amine mixture to reach a 250-fold molar excess relative to lectin. The reaction was incubated at room temperature in the dark for 2 hours. For both two strategies, after incubation, the reaction mixture was transferred to a centrifugal filter preloaded with cold lectin dilution buffer and washed three times to remove excess TFP-DBCO. Azide-labeled lectin CODEX DNA barcodes **(Table S3)** were then dissolved in lectin dilution buffer to 200 μM and added to the recovered DBCO-labeled lectins at a 5-fold molar excess relative to lectin, followed by gentle mixing. The click reaction was incubated at 4 °C for at least 12 hours.

After barcode conjugation, the reaction mixture was washed three times with cold labeled lectin storage buffer (10 mM Tris, 150 mM NaCl, 0.1 mM CaCl₂, and 0.1% sodium azide, pH 8.0) using 10 kDa or 30 kDa MWCO (Sigma Aldrich, UFC503096) centrifugal filter accordingly. After the final wash, the centrifugal filter was inverted into a new collection tube and centrifuged at 1,000 × g for 2 minutes. The recovered product was diluted to 9 μM in labeled lectin storage buffer and stored at 4°C for at least 24 hours to allow the azide-containing buffer to quench residual unreacted DBCO groups on the lectin. Conjugation efficiency was assessed by loading 1 μL of the labeled lectin onto an SDS-PAGE gel followed by staining with Oriole Fluorescent Dye (Bio-Rad, 1610495).

### Preparation of DNA-barcoded antibodies for CODEX-GP

First, the 50 kDa MWCO centrifugal filter column (Sigma Aldrich, UFC505096) was pre-equilibrated with PBST (1× DPBS with 0.1% Tween-20) to reduce nonspecific antibody binding. Antibodies supplied in PBS without additional stabilizers or additives were used directly; otherwise, the antibodies were subjected to three rounds of buffer exchange with DPBS. Next, 100 μg of each antibody was incubated with 360 μL of freshly prepared 2.5 mM Bond-Breaker TCEP Solution (Thermo Fisher Scientific, 77720). After gentle mixing, the reaction was incubated at room temperature for 30 minutes. To quench the reduction reaction and remove residual TCEP, the partially reduced antibody was washed twice with Buffer C. Buffer C contained 1 mM Tris (pH 7.0), 1 mM Tris (pH 7.5), 150 mM NaCl, 1 mM EDTA (pH 8.0), and 0.02% sodium azide in water. Afterward, 200 μg of each activated maleimide-conjugated DNA oligo **(Table S3)** was dissolved in freshly prepared high-salt Buffer C (250 mM NaCl in Buffer C) and immediately added to the partially reduced antibody solution. The conjugation reaction was then incubated at room temperature for 2 hours. After conjugation, the antibody conjugates were washed three times with high-salt PBS (1× DPBS, 900 mM NaCl, and 0.02% sodium azide in water) to remove unreacted maleimide-modified oligos. The DNA-barcoded antibodies were subsequently resuspended in 200 μL CODEX antibody stabilizer solution (stock antibody stabilizer supplemented with 495 mM NaCl and 5 mM EDTA at pH 8.0) and stored at 4 °C until use.

### Multiplex immunofluorescence imaging (CODEX) of glycans and proteins

Tissue deparaffinization and decrosslinking was performed as described for DBiT-GPT. For permeabilization, the tissue section was covered with 500 μL of 0.5% Triton X-100 in PBS and incubated at room temperature for 20 minutes, followed by three washes with DPBS. The slide was then incubated in 1× Carbo-Free Blocking Solution for 30 min at room temperature and washed three times with DPBS. For lectins staining, slide was incubated with a mixture of DNA-conjugated lectins in 1× Carbo-Free Blocking Solution for 1 hour at room temperature. The initial lectins staining concentration was 90 nM, as recommended by the manufacturer, and the staining volume was further optimized as summarized in **Table S3**.

After lectin incubation, the slide was washed three times for 5 minutes each with DPBS containing 0.1 mM CaCl_2_ under gentle agitation. Tissue-bound lectins were then fixed with 1% PFA for 5 min at room temperature, and the fixation reaction was quenched with 1.25 M lysine in DPBS. To block excess glycan-binding sites on tissue-bound lectins and avoid potential interactions with glycans on subsequently applied antibodies, slide was incubated on ice with the corresponding sugar mixtures **(Table S3)** for a total of 20 minutes, with the sugar mixture replaced once after the first 10 min. The slide was then washed three times for 5 minutes each with DPBS containing 0.1 mM CaCl_2_ under gentle agitation.

DNA-barcoded antibodies were subsequently applied for protein imaging. The initial antibody staining concentration was based on a 1:100 dilution in antibody cocktail incubation buffer supplemented with 0.1 mM CaCl2, according to the manufacturer’s recommendation, and the final staining volume was further optimized as listed in **Table S3**. Antibody cocktail staining was carried out for 3 hours at room temperature in a humidified chamber. Following antibody incubation, the tissue section was post-fixed with 1.6% PFA for 10 minutes. Finally, PhenoCycler-Fusion imaging was performed as previously described.^65^ Reporter sequences and CODEX imaging cycles are provided in **Table S3**.

### DBiT-GPT data pre-processing

For cDNA libraries derived from lectin- and antibody-associated DNA tags (LADTs), the raw FASTQ file of Read 2, which contained the UMI as well as barcode A and barcode B sequences, was first reformatted into the standard input format required by ST Pipeline version 1.8.2.^66^ LADTs-derived UMI counts were then quantified at each spatial pixel using CITE-seq-Count version 1.4.5^67^ with default settings. The resulting combined glycan and protein expression matrix contained the spatial coordinates (barcode A × barcode B) together with the corresponding glycan and protein expression levels.

For cDNA libraries derived from mRNAs, the raw FASTQ file of Read 2 was reformatted in the same manner as for LADTs-derived cDNAs. Read 1 was then aligned to the human reference genome (GRCh38) using STAR version 2.7.11b^68^ with the recommended parameters of ST Pipeline. The resulting coding gene expression matrix contained spatial coordinates (barcode A × barcode B) and corresponding gene expression levels.

Because the in situ polyadenylation strategy applied in this study enabled capture of non-coding RNAs, a whole-transcriptome matrix was additionally generated using the parameters recommended in ASTRO v 1.1.^69^ This matrix included both coding and non-coding RNAs (ncRNAs) and distinguished intronic from exonic reads, thereby increasing overall read capture and providing greater depth for RNA biology analysis in FFPE samples.

Finally, ENSEMBL IDs were converted to gene symbols for downstream analysis. Matrix entries corresponding to pixel positions without tissue were excluded. Missing pixels were inferred from neighboring data to facilitate clustering analysis across the entire mapped area.

### Unsupervised clustering analysis of DBiT-GPT data

Spatial glycan, protein, and transcript expression analyses were performed using Seurat version 5.3.0.^70^ First, raw UMI count matrices for glycan, protein, and transcript features were normalized separately for each modality across all spatial pixels using SCTransform (SCT) normalization.^71^ Linear dimensional reduction was performed using the “RunPCA” function. The number of principal components (PCs) retained for downstream analysis was determined based on the “ElbowPlot”, which visualizes the standard deviation of each PC. The selected dimensions capturing the majority of the variation in the data. Graph-based clustering was performed using the “FindNeighbors” and “FindClusters” functions. Spatial heterogeneity was visualized with “RunUMAP”, and differentially expressed glycans, proteins, and transcripts defining each cluster were identified using the “FindMarkers” function through pairwise comparisons between pixel groups.

### Correlation analysis between spatial glycans and proteins

Spatial glycan and protein count matrices were independently normalized using SCTransform. Pairwise Pearson correlations were computed between LDTs and ADTs signals pixel-by-pixel, excluding pixels with zero counts in both features. *P* values were corrected for multiple comparisons using the Benjamini–Hochberg method. These analyses were interpreted as spatial co-variation between lectin-binding motifs and protein-marker abundance, not as evidence that the corresponding protein marker carries the detected glycan motif.

### Glyco-code definition

Glyco-codes were defined as composite lectin-derived glycan signatures associated with specific histopathological regions. Glycan LDTs were normalized using SCTransform, after which region-enriched LDTs were selected for each glyco-code and z-score transformed across all spatial pixels to place individual LDTs on a comparable scale. A glyco-code score was calculated for each pixel as the average z-scored signal of all selected LDTs within that glyco-code. The resulting scores were mapped back to DBiT-GPT spatial coordinates and visualized with color scales clipped at the 5th and 95th percentiles.

### Spatial transcriptomic pseudotime analysis

Pseudotime analysis of DBiT-GPT spatial transcriptomic data was performed using Monocle 3.^72^ After preprocessing and UMAP dimensionality reduction, relevant cell populations were selected for trajectory inference. A principal graph was then learned in the reduced-dimensional space using the learn_graph function, and cells were ordered in pseudotime with the order_cells function by selecting root nodes enriched for the biologically earliest cell state in each sample. Trajectory-associated genes were identified using the graph_test function, and significant genes were ranked by Moran’s I.

### Gene set enrichment analysis with GlycoEnzOnto pathway collection

To assess differential glycosylation enzyme pathway activity between clusters, gene set enrichment analysis (GSEA) was performed using the fgsea package version 1.26.0^73^ in R version 4.4.0. A signed ranking metric, defined as sign(avg_log_2_FC) × (−log_10_(*P*adj)), was computed for each differentially expressed gene to jointly capture the direction and statistical significance of expression changes. Genes were ranked in descending order by this metric and used as input for fgsea run against the GlycoEnzOnto pathway collection,^74^ a curated GMT database of glycosylation-related enzymatic pathways. Pathways with Benjamini–Hochberg adjusted *P* < 0.05 were considered significantly enriched.

### Liver zonation score

Liver zonation scores were calculated using curated zone-specific hepatocyte marker genes: Zone 1 (*HAL*, *CPS1*, *ASS1*, *OTC*, *ARG1*), Zone 2 (*HSD17B13*, *CYP8B1*, *HPD*, *PON1*), and Zone 3 (*GLUL*, *CYP2E1*, *CYP3A4*, *CYP3A5*, *AXIN2*). Module scores were computed on the SCT-normalized transcriptomic assay using Seurat AddModuleScore and mapped back to spatial coordinates using SpatialFeaturePlot, with color scales clipped at the 5th and 95th percentiles.

### Ingenuity Pathway Analysis

Ingenuity Pathway Analysis (IPA, QIAGEN) was performed on differentially expressed transcripts defining individual clusters to identify associated canonical pathways and interaction networks. Predicted pathway activity was assessed using the IPA activation z-score, with positive and negative values indicating activation and inhibition, respectively; absolute z-scores ≥2 was considered significant. Pathway enrichment significance was determined by right-tailed Fisher’s exact test. Networks were generated using the heuristic graph algorithm implemented in IPA.

### Integrated analysis of spatial and scRNA-seq and snRNA-seq datasets

Public scRNA-seq and snRNA-seq datasets (GSE149614^75^ and GSE202379^76^) were obtained from the Gene Expression Omnibus (GEO) and integrated with our spatial datasets using Seurat. Pixels with fewer than 1,000 read counts were excluded. The data were then normalized using SCTransform. Principal component analysis (PCA) was performed on the SCTransform-normalized data, followed by Harmony correction on the first 25 principal components using data source as the batch variable. UMAP was computed from the Harmony embedding, and graph-based clustering was performed on the first 15 Harmony dimensions at a resolution of 0.2. For tissue type annotation, clustering was performed independently for each spatial dataset using a similar analytical workflow, and clusters were manually annotated based on pathological image features together with gene expression profiles.

Liver tissue-specific markers were identified based on tissue type annotation using the wilcoxauc function from the presto package version 1.0.0. Markers were ranked by AUC score, and the top-ranking markers were selected. Mitochondrial genes, ribosomal genes, small RNAs, antisense transcripts and poorly annotated loci were excluded before marker selection. Average normalized expression was then calculated for each cluster using the AverageExpression function, scaled by gene-wise Z-score transformation and visualized as heatmaps using pheatmap package version 1.0.12. In addition, canonical marker genes for liver fibrosis, steatosis and tumor states were visualized separately using the same procedure.

### Cell segmentation of CODEX images

Multiplex fluorescence CODEX images were segmented at the single-cell level using Mesmer by DeepCell.^77^ Prior to segmentation, the nuclear DAPI channel was isolated and preprocessed using contrast-limited adaptive histogram equalization (CLAHE) with a tile size of 599 px, a slope of 10, and 8-bit histogram bins to calculate the clip limit. This was followed by box filter smoothing (3 px radius) and unsharp masking (radius 7, amount 2). A pseudo-membrane channel was generated by taking the average of the mean and maximum intensity projections of the remaining marker channels and subtracting the preprocessed DAPI channel. The preprocessed DAPI channel was utilized exclusively for segmentation purposes to ensure that local intensity variations and overall texture did not influence cell boundary accuracy.

### Adjacent IHC quality control alignment and correlation

To validate the simultaneous application of glycan and protein markers, ground-truth IHC images were spatially aligned to the CODEX coordinates. The immediately adjacent FFPE sections from the 3 HCC subtype samples used for CODEX imaging were stained with anti-HepPar-1 antibodies and detected through secondary DAB-functionalized antibodies. Briefly, high-resolution OME-TIFFs were loaded and subjected to automated tissue detection utilizing morphological operations, including closing, opening, and hole-filling, so that a rigid-body scaling Intersection over Union (IoU) co-registration can be performed on the tissue boundaries for both IHC and CODEX fluorescence images, then a Fast Fourier Transform (FFT)-based phase cross-correlation fine-tuned the registration. IHC images were acquired at 40× magnification and rescaled to match the spatial resolution of the 20× CODEX images. Co-registration was performed using rigid-body alignment of tissue outlines using VISTAmap^78^ followed by fine adjustment based on texture-based 2D cross-entropy between the grayscale hematoxylin channel from the IHC image and the DAPI channel from the CODEX image. Co-registered IHC and CODEX signals were then compared using Spearman correlation coefficients, Manders M2 scores, and Dice scores. Differentiation heatmaps shown in Supplementary Figure S13 were calculated using the min-max normalized 99.5^th^ percentile Otsu thresholding and subtractive color mixing as follows:

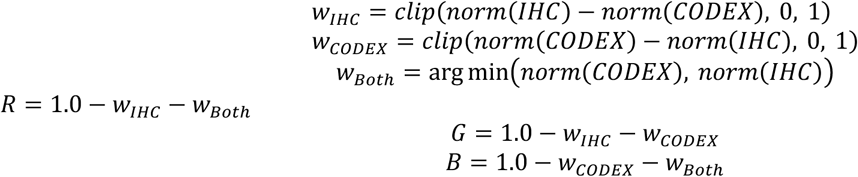

### Cell-cell proximity-based spatial protein–glycan interactome analysis

Spatial coordinates (centroid X, centroid Y) from each tissue core were used to construct a physical proximity graph. Cell interactions were visualized as circular chord diagrams using the Python package pyCirclize version 1.10.1, in which arc widths encode the number of participating cell types and ribbon widths encode adjacency events. To characterize the spatial co-occurrence patterns between protein-defined and glycan-defined cell populations, a cross-modality spatial interactome was constructed. For each core, a k-dimensional tree was constructed from cell centroids using cKDTree from SciPy, and all cell pairs within a Euclidean contact radius of 30 coordinate units were identified as putative cell–cell contacts. Observed contact counts were tallied into interaction matrices separately for protein-defined clusters (homotypic), glycan-defined clusters (homotypic), and cross-modality pairs (heterotypic). Each matrix was converted to a Log_2_ odds enrichment score comparing observed contacts to the frequency expected under spatial randomness. Interaction pairs exceeding a Log_2_ odds threshold of 0.5 were retained for visualization. The cross-modality enrichment matrix was rendered as a network graph in which edge weights encode the Log_2_ odds enrichment score, with ribbons below threshold suppressed to highlight the strongest spatial co-associations. Separate heatmaps of all three log_2_ odds matrices were generated using the clustermap function in seaborn^79^ to provide a comprehensive view of the spatial interactome.

### Clustering of CODEX images

Multiplex fluorescence CODEX images were segmented as described above. Cell-mean intensity values for each channel were winsorized at the bottom 10th percentile to zero, log1p transformed, and z-score transformed prior to further analysis. GPU-accelerated K-Means clustering (RAPIDS cuML) was applied to define cellular populations, which were subsequently visualized using Uniform Manifold Approximation and Projection (UMAP). Cell types were annotated manually and assigned to unsupervised clusters.

### Quantification of CODEX marker contribution to spatial phenotypes

To identify lectin and protein markers that most strongly contributed to spatial phenotype identity, a random forest classifier was trained to predict cluster membership using single-cell marker intensities as input features. The classifier was implemented using the RandomForestClassifier function in scikit-learn with 100 trees and a maximum depth of 12. Gini-based feature importance scores were extracted and ranked separately for cell-type cluster labels, using all cells, and for spatial path or trajectory labels, using cells annotated by tissue region. To quantify the information-theoretic contribution of each marker to cluster resolution, Shannon entropy was computed from conditional cluster-assignment probability distributions derived from the clustering confusion matrix. For each cell, the entropy of the glycan-cluster distribution conditioned on protein-cluster assignment, and vice versa, was calculated to assess modality-specific divergence. Relative entropy, measured as Kullback–Leibler divergence, was used to estimate the additional information gained by incorporating glycan markers beyond protein markers alone, thereby evaluating whether glycans encode biological variation not captured by protein features.

### Clustering comparison and granularity agreement

To evaluate the independent and complementary contributions of glycan markers, clustering performance was assessed across a range of k values using multiple validation metrics, including the Calinski-Harabasz score, silhouette score, adjusted Rand index, and Davies-Bouldin index. Protein-only clustering was used as the reference partition. Glycan-only and combined protein–glycan clusters were then aligned to the protein-only reference to enable consistent visualization across UMAP embeddings and spatial maps. Cluster alignment was performed using a hybrid coloring strategy that combined Hungarian matching with spatial parent-tracing, preserving shared lineage relationships while highlighting additional substructure. Granularity was assessed by comparing whether cells assigned to the same protein-defined cluster were further subdivided in glycan-only or combined clustering, and vice versa. A clustering result was considered to have higher granularity when a cluster in one modality contained multiple distinct clusters defined by the other modality.

### Subcellular glycans distribution analysis

Raw CODEX images and segmentation masks were paired and allocated a global prototype-cell budget of 1,000,000 unique cells across slides and then carried those sampled cells through compartment-aware feature extraction, broad cell-type conditioning, and conditional generative modeling. Before compartment inference, each sampled prototype cell mask was dilated by 10% to avoid clipping membrane-proximal signal. For each prototype cell, nuclear, cytoplasmic, membrane, and concentric radial-ring regions were inferred from the binary whole-cell mask using a Euclidean distance transform. Geometric features, including area, eccentricity, major and minor axis lengths, and orientation, were computed on the GPU, whereas solidity and Haralick texture features were computed crop by crop on the CPU. Subcellular distribution was represented by relative localization rather than raw abundance. Accordingly, raw compartment and radial-ring mean intensities were normalized to the corresponding whole-cell mean for each marker. Because segmented cell areas were left-skewed, cells within the bottom 10% of the area distribution were excluded. The remaining cells were centroid-aligned, stored in Zarr format, and 90 percent of remaining cells were used to train the conditional variational autoencoder (cVAE), with a binary cell-mask channel concatenated to the glycan channels as part of the encoder input.

The conditioning variable for the cVAE was the broad antibody-defined cell type annotation. The model input consisted of multi-channel glycan image crops concatenated with one binary cell-mask channel. The encoder contained four convolutional blocks that progressively downsampled the input from 64 × 64 to 32 × 32, 16 × 16, 8 × 8, and 4 × 4 feature maps. A learned condition embedding with dimension 16 was incorporated into a 32-dimensional latent space, followed by a mirrored transposed-convolution decoder that reconstructed the multi-channel glycan image. Reconstruction loss was mask-weighted so that the model retained the full square crop while assigning greater weight to the cell body than to the background. The model was trained with a batch size of 64 for 8 epochs using a learning rate of 1 × 10^-3^, a latent dimension of 32, and a β value of 1 × 10^-3^.

### Pseudo-bulk glycans expression by liver disease

Single-cell lectin-derived glycan signals from CODEX-GP TMA cores were aggregated to generate pseudo-bulk glycan-class profiles for each disease group. For each tissue core, the mean raw intensity of each lectin-derived glycan signal was computed across all cells, yielding a core-level lectin signal matrix. Lectins were then grouped into seven biologically defined glycan-binding classes based on their major binding specificities: mannose-binding lectins (ConA, LCA, PSA), fucose-binding lectins (AAL, UEA I, LTL), sialic acid-binding lectins (SNA, MAL-I, MAL-II), GlcNAc-binding lectins (WGA, LEL), galactose-binding lectins (PNA, ECL, BPL, GSL-I), GalNAc-binding lectins (SBA, VVL, DBA, WFA, MPL, Jacalin), and complex N-glycan-binding lectins (PHA-E, PHA-L). To place all lectin signals on a common scale irrespective of absolute staining intensity, each core-level lectin mean was normalized by z-score transformation relative to the Normal disease group by subtracting the cross-core mean of Normal cores and dividing by the standard deviation of Normal cores for that lectin. A class-level glycan score was then calculated as the mean normalized z-score across all lectins assigned to that glycan-binding class. Cores were assigned to one of eight harmonized disease groups (Normal, Steatosis/FLD, Hepatitis, Fibrosis, Cirrhosis, HCC, CCA, and clear cell HCC) using a standardized pathology-diagnosis crosswalk; groups with fewer than three cores were excluded. Class-level glycan scores were displayed as grouped box plots with individual core-level data points overlaid, with a dashed reference line at zero representing the Normal baseline and a red line connecting group means across disease stages to visualize disease-associated trends.

### Multimodal classification of liver disease

To benchmark the discriminative value of lectin-derived glycan profiles relative to, and in combination with, protein profiles, we applied a multimodel classification framework at the tissue-core level. Core-level feature vectors were generated by aggregating single-cell marker intensities into per-core summary statistics, including the mean intensity and positive-cell fraction for each marker. Thus, each TMA core, rather than each cell, represented one independent sample for model training and evaluation. Cross-validation splits were performed at the core level, so single-cell measurements and spatial-neighborhood features derived from the same core were never divided between training and held-out folds. This design reduces leakage caused by intra-core spatial autocorrelation among neighboring cells. Three marker panels were evaluated in parallel: antibody markers alone, lectin-derived glycan markers alone, and combined antibody and lectin markers. Four classifier architectures were benchmarked: elastic-net logistic regression, ExtraTrees, XGBoost, and a feedforward neural network implemented as a multilayer perceptron (MLP) in PyTorch. All classifiers were evaluated using core-level stratified k-fold cross-validation with StratifiedKFold in scikit-learn to preserve class balance across splits. Model performance was assessed on held-out fold predictions using macro-averaged and weighted one-versus-rest AUROC, accuracy, and average precision. Feature importance scores were extracted from tree-based ExtraTrees models to rank markers by discriminative contribution. As a spatial extension, tabular core-level features were further augmented with spatial autocorrelation statistics derived from the cell proximity graph.

### PCA-based analysis of glycan signatures and covariate-adjusted disease/stage associations

Class-level glycan scores, defined as the mean log1p-transformed lectin intensity for each glycan-binding class within each core, were used to construct a PCA space. Scores were standardized to zero mean and unit variance using StandardScaler before PCA decomposition. Clinical metadata, including disease group, sex, age, fibrosis stage, and histological grade, were reserved exclusively for downstream statistical modeling and were not used to derive the PCA axes. To distinguish disease-associated variation from covariate-associated variation in PC scores, a nested ordinary least-squares (OLS) F-test framework was applied. For each PC, a reduced model containing only nuisance covariates, including age and sex, was compared with a full model that additionally included the biological variable of interest, such as disease group, stage, or grade. The incremental F-statistic from this nested comparison was used to derive a covariate-adjusted *P* value for each biological factor. Multiple testing was controlled using both Benjamini-Hochberg false discovery rate (FDR) q values and Bonferroni-adjusted *P* values. Within individual disease groups, analogous nested F-tests were performed to evaluate stage-only, grade-only, and combined stage-plus-grade effects on PC scores, with additional confounding tests used to assess whether age was associated with stage or grade. Bidirectional BIC-based stepwise model selection was applied to identify the minimal model that best explained variation in each PC score. Partial R^2^ was used to quantify the unique contribution of each factor after adjustment for all other variables.

### Lectin treatment and clonogenic assay

Freshly prepared lectins were dissolved in lectin dilution buffer and filtered through 0.22 μm filter (Sigma Aldrich, SLGSR33SS) before use. Huh7 cells in good condition were dissociated with trypsin (Thermo Fisher Scientific, 25200056), resuspended in complete culture medium, counted, and seeded into 6-well plates at a density of 400 cells per well. Lectins were added to the culture medium at a final concentration of 10 μg/mL. Cells were cultured for 14 days, or until most colonies contained more than 50 cells, with the culture medium replaced every 3 days and fresh lectins added at each medium change. After colony formation, cells were gently washed with PBS, fixed with 1 mL of 4% PFA for 60 minutes, and washed again with PBS. Colonies were then stained with 1 mL of crystal violet solution (Sigma Aldrich, V5265) for 20 minutes. Colony formation was quantified using the ColonyArea plugin in ImageJ,^80^ which automatically measures colony growth in clonogenic assays.

### siRNA transfection, knockdown validation, and clonogenic assay

For siRNA-mediated knockdown, Huh7 cells were seeded into 6-well plates at a density of 6 × 10^5^ cells per well. After overnight attachment, cells were transfected with 25 pmol siRNA per well using RNAiMAX transfection reagent (Thermo Fisher Scientific, 13778150) according to the manufacturer’s instructions. A non-targeting siRNA was used as the negative control, and the siRNA–transfection reagent complexes were added dropwise to each well. After 72 hours of transfection, knockdown efficiency was evaluated by Quantitative real-time PCR (RT-qPCR). siRNAs with validated knockdown efficiency were then used for subsequent clonogenic assays. Briefly, siRNA-transfected cells were seeded into 6-well plates at a density of 400 cells per well for colony formation. Cells were cultured for 10-14 days until visible colonies formed, defined as colonies containing more than 50 cells, with the culture medium replaced every 3 days. Colony fixation, crystal violet staining, and quantification were performed as described above for the lectin treatment.

### Statistical analysis

Statistical analyses were performed using R version 4.4.0 (R Foundation for Statistical Computing, Austria) and GraphPad Prism version 10.6.1 (GraphPad Software, USA). The specific statistical tests used are indicated where applicable.

