## Supplementary Information for "Spatial Glyco-Codes Define Human Liver Pathology and Progression"

**Table S1 Chemicals and reagents**

| Chemicals and reagents | Catalog # | Vendor |
|---|---|---|
| Maxima H Minus Reverse Transcriptase | EP0753 | Thermo Fisher Scientific |
| KAPA HiFi HotStart ReadyMix (2X) | KK2602 | Kapa Biosystems |
| T4 DNA Ligase | M0202L | New England Biolabs |
| D5000 ScreenTape | 5067-5588 | Agilent Technologies |
| D5000 ladder reagents | 5067-5589 | Agilent Technologies |
| dNTP Mix | R0192 | Thermo Fisher Scientific |
| RNAse Inhibitor | Y9240L | QIAGEN |
| SUPERase·In™ RNase Inhibitor (20 U/μL) | AM2694 | Thermo Fisher Scientific |
| Fisher BioReagents Formaldehyde (37% by Weight/Molecular Biology) | BP531-25 | Fisher Scientific |
| NEBuffer™ r3.1 | B6003S | New England Biolabs |
| *E.coli* Poly (A) Polymerase | M0276L | New England Biolabs |
| Cell Staining Buffer | 420201 | BioLegend |
| EvaGreen® Plus Dye, 20X | 31077-T | Biotium |
| SEQuoia RiboDepletion Kit, 24 reactions | 17006487 | Bio-Rad |
| Dynabeads™ MyOne™ Streptavidin C1 | 65001 | Thermo Fisher Scientific |
| Proteinase K, recombinant, PCR grade | EO0491 | Thermo Fisher Scientific |
| SPRIselect DNA Size Selection Reagent, 60 mL | B23318 | Beckman Coulter |
| Streptavidin Conjugation Kit - Lightning-Link® | ab102921 | abcam |
| Antigen Retrieval Buffer (100X Tris-EDTA Buffer, pH 9.0) | ab93684 | abcam |
| EZ-Link™ TFP Ester-PEG4-DBCO | C20043 | Thermo Fisher Scientific |
| Oriole Fluorescent Gel Stain | 1610495 | Bio-Rad |
| 10mM Tris with 0.1% Sodium Azide, pH 8.0 | 40121289 | bioWORLD |
| 1M $CaCl_2$ | 21115 | Sigma Aldrich |
| 10X DPBS | 14200075 | Thermo Fisher Scientific |
| TRIS, 1.0M buffer soln., pH 7.0 | J62657.K2 | Thermo Fisher Scientific |
| TRIS, 1.0M buffer soln., pH 7.5 | J60636.K2 | Thermo Fisher Scientific |
| EDTA (0.5 M), pH 8.0, RNase-free | AM9260G | Thermo Fisher Scientific |
| Tween™ 20 Surfact-Amps™ Detergent Solution | 85113 | Thermo Fisher Scientific |
| Bond-Breaker™ TCEP Solution, Neutral pH | 77720 | Thermo Fisher Scientific |
| 1% Sodium Azide | 786-299 | G-Biosciences |
| 5M NaCl | 71386 | Sigma Aldrich |
| Boca Scientific Inc Antibody Stabilizer PBS Base, 50ml | NC0436689 | Fisher Scientific |
| Sample Kit for PhenoCycler-Fusion | 7000017 | Quanterix |
| 10X Buffer Kit for PhenoCycler-Fusion | 7000019 | Quanterix |
| Assay Reagent for PhenoCycler | 7000002 | Quanterix |
| Nuclear Stain for PhenoCycler | 7000003 | Quanterix |

**A** Clinicopathological information of the 12 patients

| Tumor ID | Background ID | Background liver disease | HCC steatosis | Block storage duration | Age | Gender |
|---|---|---|---|---|---|---|
| HCC-1T | HCC-1N | HCV | ~70% | 8 years | 67 | Male |
| HCC-2T | HCC-2N | HCV | ~30% | 8 years | 66 | Male |
| HCC-3T | / | HBV | ~10% | 8 years | 36 | Female |
| HCC-4T | / | Unknown | <5% | 8 years | 75 | Female |
| HCC-5T | HCC-5N | HCV | <5% | 8 years | 58 | Male |
| HCC-6T | HCC-6N | HBV | ~30% | 8 years | 37 | Male |
| MASH-HCC-1T | / | MASH | ~60% | 7 years | 78 | Female |
| MASH-HCC-2T | / | MASH | ~80% | 6 years | 65 | Male |
| MASH-HCC-3T | / | MASH | ~40% | 5 years | 65 | Male |
| MASH-HCC-4T | / | MASH | ~65% | 3 years | 71 | Male |
| MASH-HCC-5T | / | MASH | ~50% | 2 years | 75 | Female |
| MASH-HCC-6T | / | MASH | ~40% | 2 years | 59 | Female |

**B** Quality control summary of DBiT-GPT across 16 blocks

| Sample ID | Glycan | | Protein | | Transcript | |
|---|---|---|---|---|---|---|
| | UMIs | Glycans | UMIs | Proteins | UMIs | Transcripts |
| HCC-1T | 9630.72 | 28.96 | 113.71 | 44.44 | 2152.76 | 400.83 |
| HCC-2T | 11377.32 | 28.96 | 1847.1 | 128.57 | 16904.89 | 3355.61 |
| HCC-3T | 13687.07 | 29.03 | 693.78 | 106.02 | 2934.27 | 727.92 |
| HCC-4T | 15837.07 | 28.88 | 1186.44 | 117.99 | 3000.62 | 765.76 |
| HCC-5T | 3816.93 | 28.35 | 320.18 | 86.24 | 17990.75 | 6236.43 |
| HCC-6T | 11386.35 | 29.02 | 110.89 | 46.51 | 4687.50 | 744.68 |
| HCC-1N | 10196.28 | 28.56 | 212.59 | 68.42 | 5221.78 | 613.04 |
| HCC-2N | 6285.67 | 26.73 | 663.21 | 101.08 | 18197.62 | 4030.26 |
| HCC-5N | 11653.58 | 28.58 | 3006.94 | 140.81 | 5213.51 | 2158.22 |
| HCC-6N | 14126.45 | 28.89 | 266.49 | 71.81 | 967.12 | 156.48 |
| MASH-HCC-1T | 3353.77 | 27.28 | 200.65 | 65.09 | 22391.87 | 4601.63 |
| MASH-HCC-2T | 8629.81 | 28.77 | 49.08 | 23.98 | 2951.41 | 1249.68 |
| MASH-HCC-3T | 4914.44 | 26.75 | 482.97 | 96.26 | 17854.69 | 4825.07 |
| MASH-HCC-4T | 4523.68 | 24.3 | 81.53 | 39.09 | 16327.24 | 3932.38 |
| MASH-HCC-5T | 8251.97 | 29.02 | 121.06 | 46.5 | 9912.26 | 4075.14 |
| MASH-HCC-6T | 7468.54 | 28.12 | 187.48 | 66.78 | 18747.28 | 6343.87 |

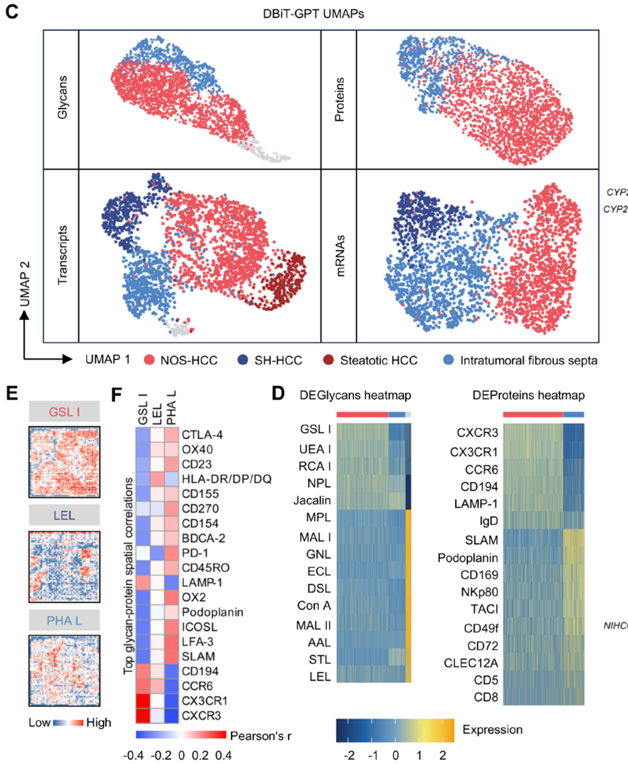


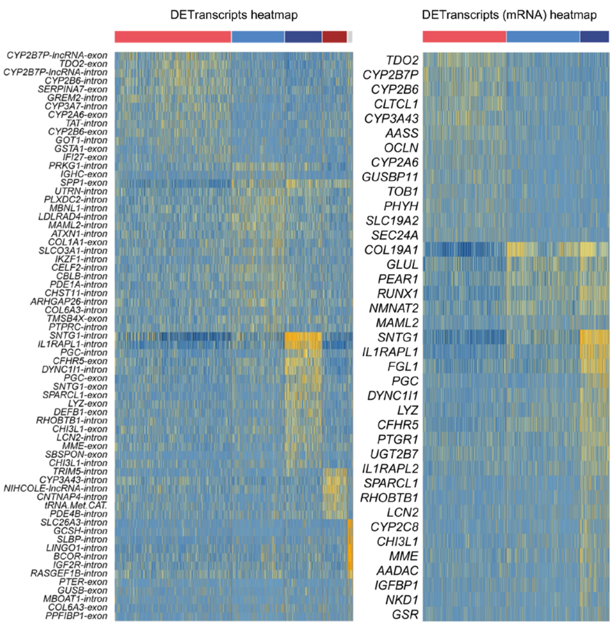


**Figure S1 Clinicopathological overview of the study cohort, DBiT-GPT quality control, and representative multi-omics clustering in sample HCC-2T, related to Figure 1**

(A) Clinicopathological information for the 12 patients included in this study, including HCC IDs, matched background sample IDs, background liver disease, degree of HCC steatosis, FFPE block storage duration, age, and gender.

(B) Quality control summary of DBiT-GPT across 16 FFPE blocks from the 12 patients included in this study. For each block, the table summarizes the average UMI counts and detected feature counts per spot for glycans, proteins, and transcripts.

(C) UMAP visualizations of glycan, protein, and transcript clusters identified in sample HCC-2T.

(D) Heatmap of the top differentially expressed glycans, proteins, transcripts, and mRNAs defining each cluster, shown from left to right.

(E–F) Spatial maps of representative cell type-associated lectin-binding glycan signals (E), and heatmap of top spatially co-localized proteins identified by pairwise Pearson correlations (F).

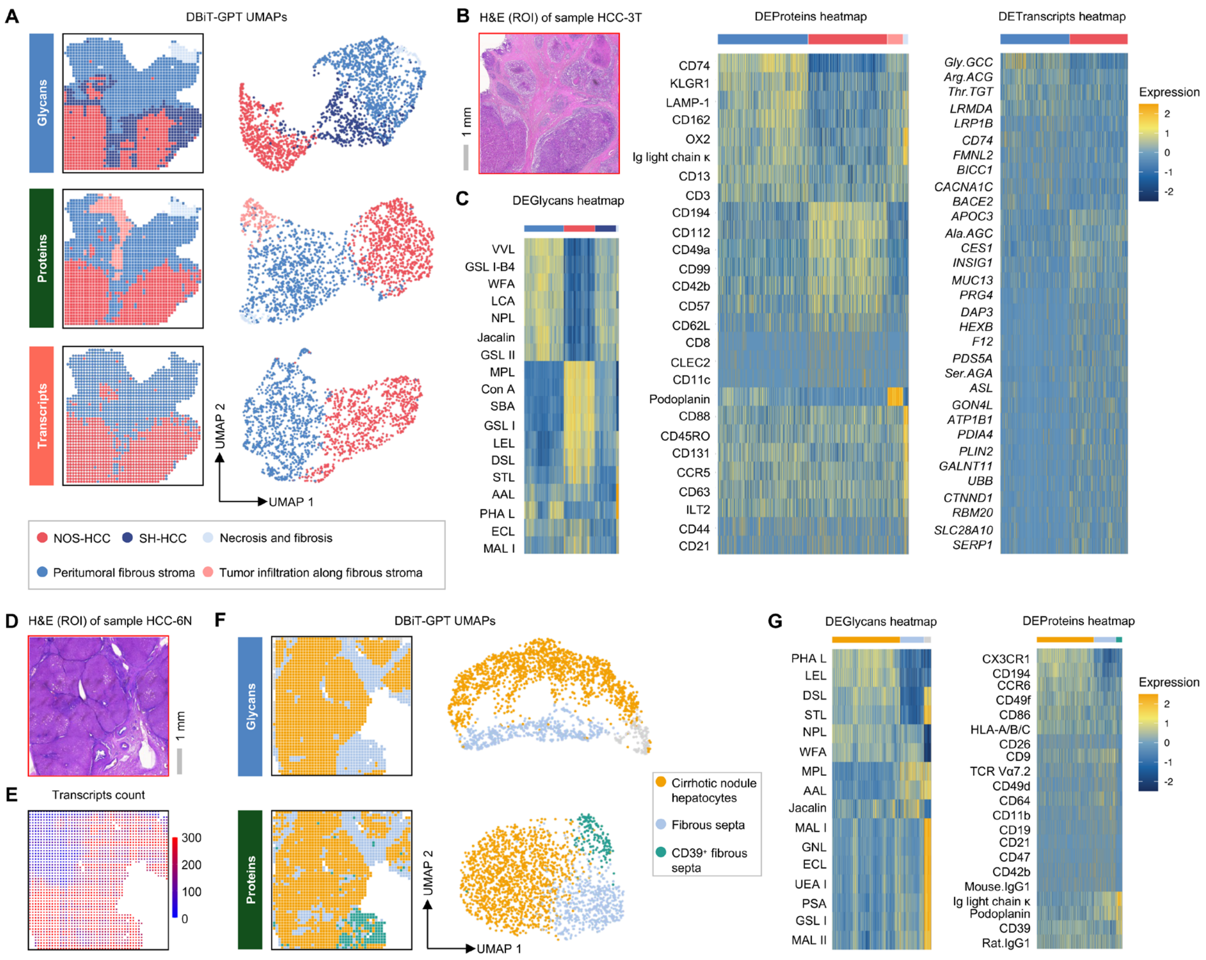


**Figure S2 Multi-omics clustering in sample HCC-3T and HCC-6N, related to Figure 1**
(A–C) Spatial clustering results of DBiT-GPT in sample HCC-3T. (A) Unsupervised clustering based on spatial glycome, proteome, and transcriptome data. (B) H&E (ROI) staining of an adjacent section. (C) Heatmap of the top differentially expressed glycans, proteins, and transcripts defining each cluster, shown from left to right.
(D–G) Spatial clustering results of DBiT-GPT in sample HCC-6N. (D) H&E (ROI) staining of an adjacent section. (E) Spatial QC map showing transcript feature counts per spot.
(F) Unsupervised clustering based on spatial glycome and proteome data. (G) Heatmap of the top differentially expressed glycans and proteins defining each cluster, shown from left to right.

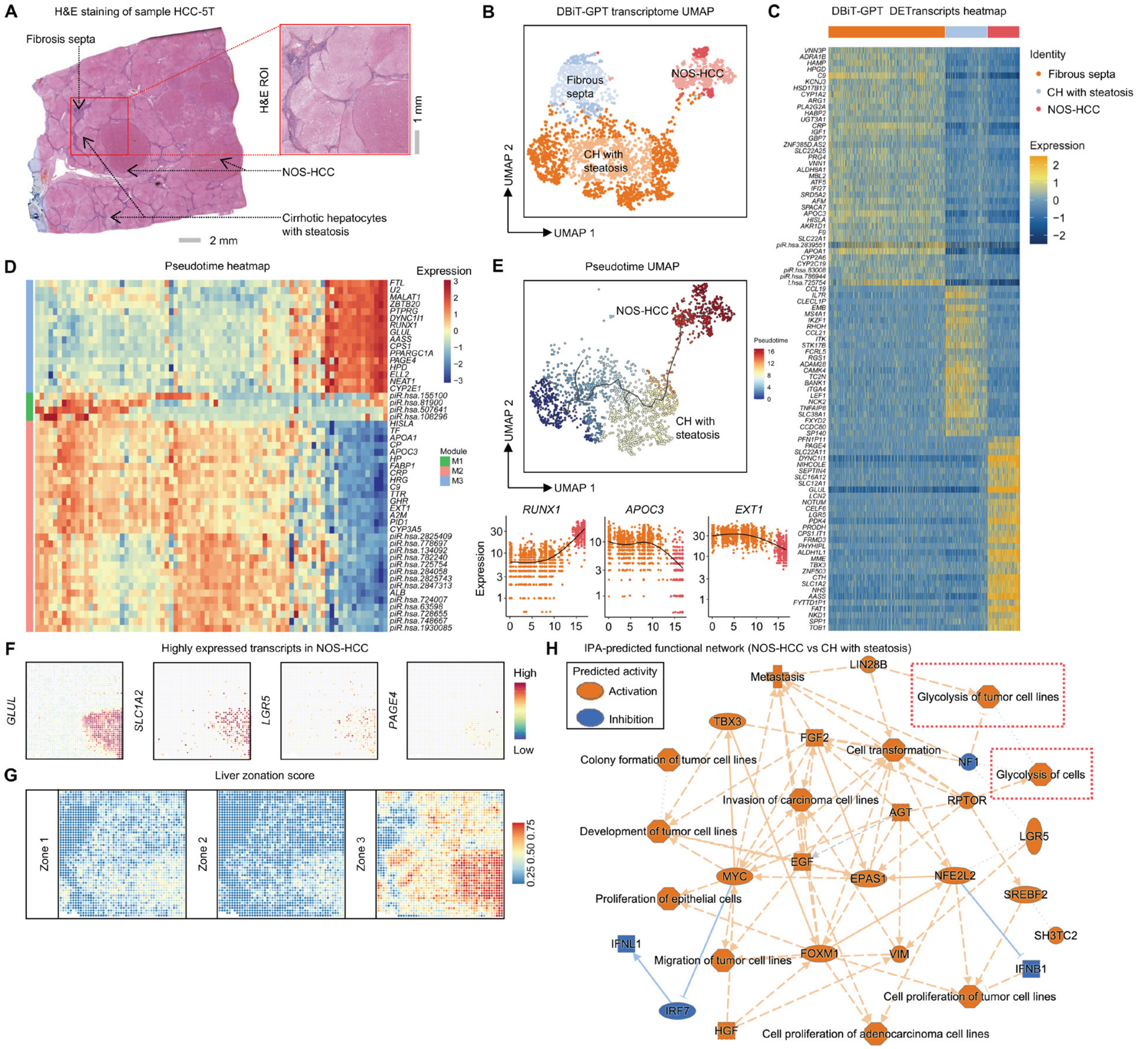


**Figure S3 DBiT-GPT profiling of sample HCC-5T, related to Figure 2**

(A) H&E staining of the full adjacent section.

(B) UMAP of unsupervised clustering based on DBiT-GPT transcriptome.

(C) Heatmap of differentially expressed transcripts (DETranscripts) identified from unsupervised clustering of DBiT-GPT transcriptome.

(D) Heatmap of transcripts dynamically regulated along pseudotime inferred from spatial transcriptome, showing distinct gene modules associated with the pseudotemporal transition.

(E) Top: pseudotime UMAP showing a trajectory from CH with steatosis to NOS-HCC. Bottom: representative transcripts dynamically changing along pseudotime.

(F) Spatial plots of highly expressed transcripts in NOS-HCC selected for *in vitro* validation.

(G) Spatial maps of liver zonation scores inferred from spatial transcriptomic annotations, showing the distributions of zone 1 (*HAL*, *CPS1*, *ASS1*, *OTC*, *ARG1*), zone 2 (*HSD17B13*, *CYP8B1*, *HPD*, *PON1*), and zone 3 (*GLUL*, *CYP2E1*, *CYP3A4*, *CYP3A5*, *AXIN2*) programs.

(H) Ingenuity Pathway Analysis (IPA)-predicted functional network comparing NOS-HCC and CH with steatosis. Input genes were filtered using *P*adj < 0.01 and $Log_2$ fold change ⩾ 1 or ⩽ -1.

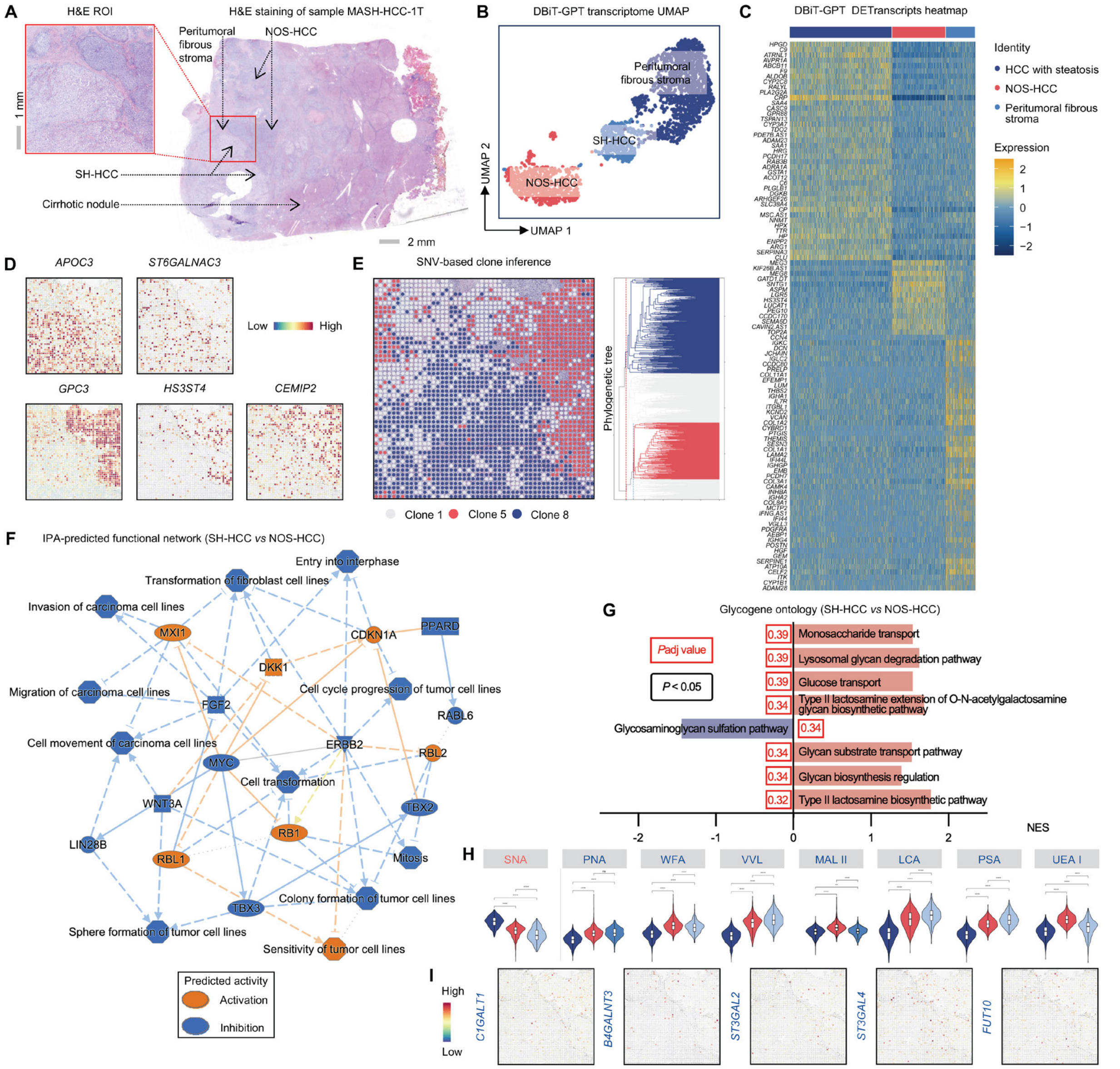


**Figure S4 DBiT-GPT profiling of sample MASH-HCC-1T, related to Figure 2**

(A) H&E staining of the full adjacent section.

(B) UMAP of unsupervised clustering based on DBiT-GPT transcriptome.

(C) Heatmap of DETranscripts identified from unsupervised clustering of DBiT-GPT transcriptome.

(D) Spatial plots of transcripts highly expressed in SH-HCC (top) and NOS-HCC (bottom).

(E) Left: inferred somatic variant patterns based on DBiT-GPT transcriptome, revealing the spatial distribution of SNV-based inferred clones. Right: phylogenetic tree of the inferred clones.

(F) IPA-predicted functional network comparing SH-HCC and NOS-HCC. Input genes were filtered using *P*adj < 0.01 and $\text{Log}_2$ fold change ⩾ 1 or ⩽ -1.

(G) Glycogene ontology analysis comparing clusters of SH-HCC and NOS-HCC. All displayed pathways had nominal *P* values < 0.05, and *P*adj values are also shown in red color.

(H) Violin plots showing the distributions of individual lectin-binding glycans across all clusters, corresponding to those shown in Figure 2T. Statistical significance across clusters was evaluated using a Kruskal-Wallis test followed by pairwise Wilcoxon rank-sum tests with Benjamini-Hochberg correction for multiple comparisons. Significance levels are indicated as ns, not significant; *, *P*adj < 0.05; **, *P*adj < 0.01; ***, *P*adj < 0.001; and ****, *P*adj < 0.0001.

(I) Spatial plots of significant glycogenes identified from the differentially expressed transcripts shown in Figure 2S.

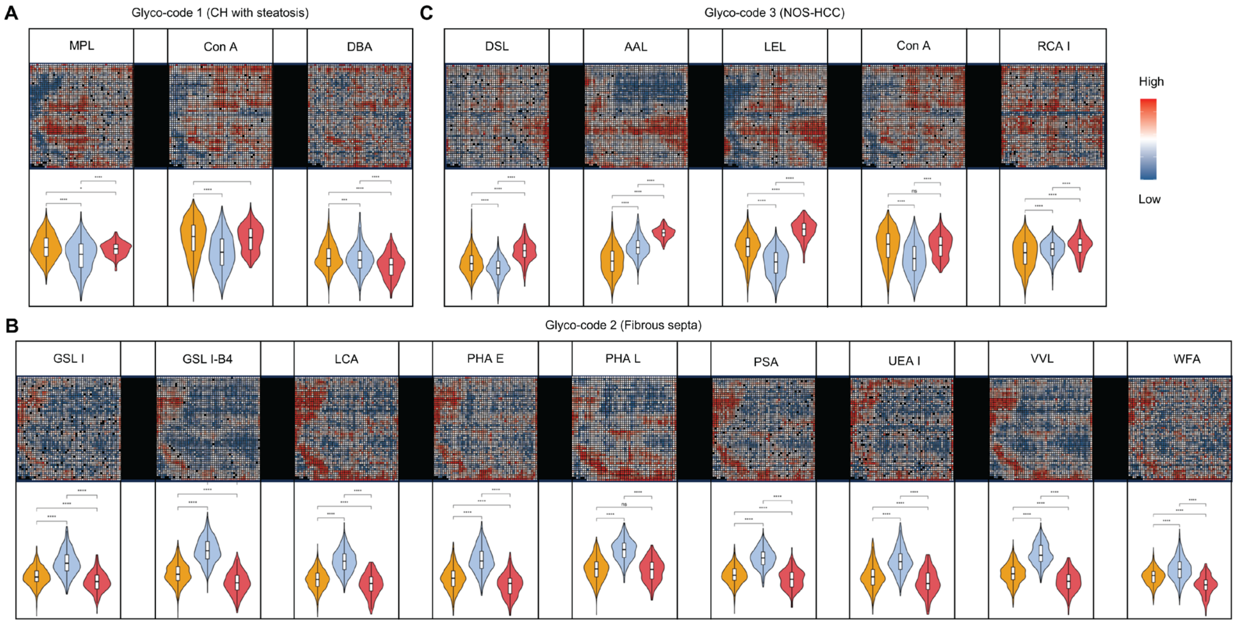


**Figure S5 Glyco-code and glycans expression of sample HCC-5T, related to Figure 2**
(A–C) Glyco-codes 1, 2, and 3 identified in HCC-5T, corresponding to CH with steatosis (A), fibrous septa (B), and NOS-HCC (C), respectively. For each glyco-code, the upper panels show the spatial distribution of individual lectin-binding glycans, and the lower panels show the corresponding violin plots with statistical comparisons across clusters. Statistical significance across clusters was evaluated using a Kruskal-Wallis test followed by pairwise Wilcoxon rank-sum tests with Benjamini-Hochberg correction for multiple comparisons. Significance levels are indicated as ns, not significant; *, *P*adj < 0.05; **, *P*adj < 0.01; ***, *P*adj < 0.001; and ****, *P*adj < 0.0001.

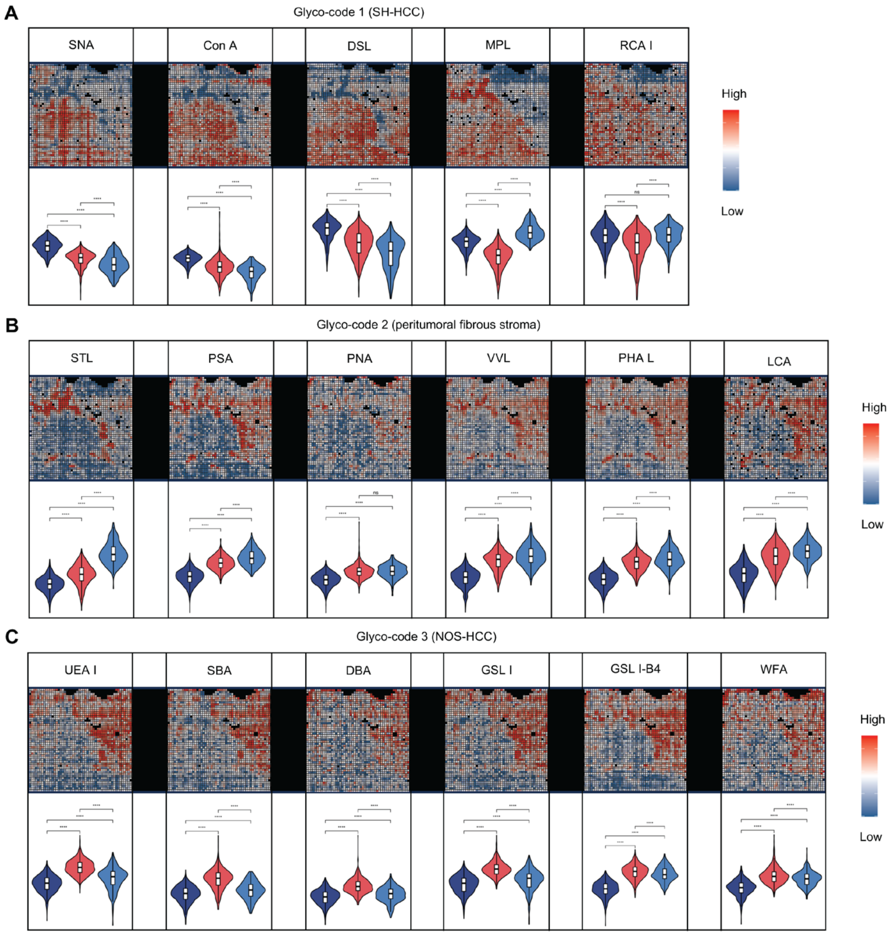


**Figure S6 Glyco-code and glycans expression of sample MASH-HCC-1T, related to Figure 2**
(A–C) Glyco-codes 1, 2, and 3 identified in MASH-HCC-1T, corresponding to SH-HCC (A), peritumoral fibrous stroma (B), and NOS-HCC (C), respectively. For each glyco-code, the upper panels show the spatial distribution of individual lectin-binding glycans, and the lower panels show the corresponding violin plots with statistical comparisons across clusters. Statistical significance across clusters was evaluated using a Kruskal-Wallis test followed by pairwise Wilcoxon rank-sum tests with Benjamini-Hochberg correction for multiple comparisons. Significance levels are indicated as ns, not significant; *, *P*adj < 0.05; **, *P*adj < 0.01; ***, *P*adj < 0.001; and ****, *P*adj < 0.0001.

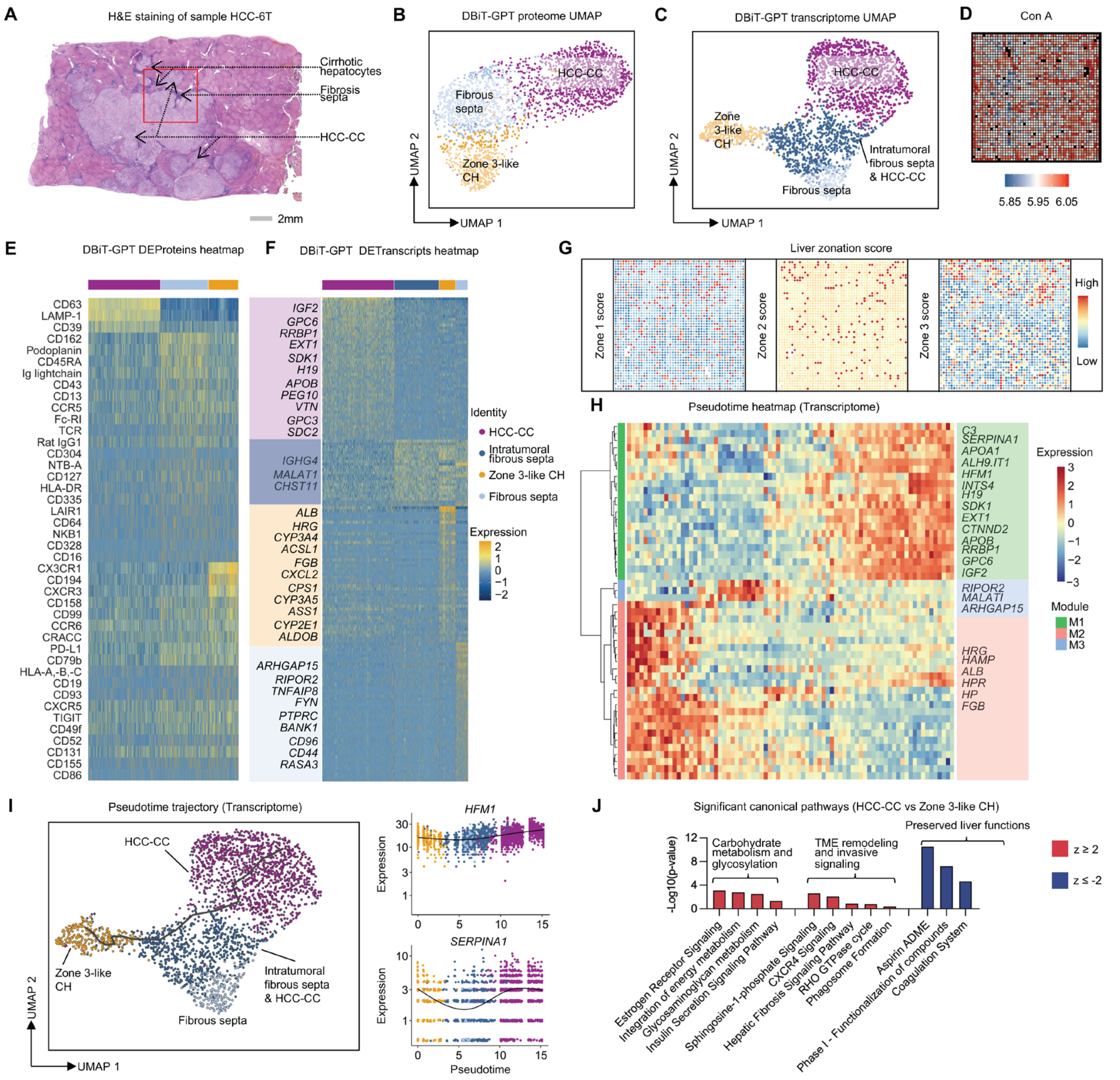


**Figure S7 DBiT-GPT profiling of sample HCC-6T, related to Figure 3**

(A) H&E staining of the full adjacent section.

(B–C) UMAPs of unsupervised clustering based on DBiT-GPT proteome (B) and transcriptome (C) data, respectively.

(D) Spatial distribution of Con A-binding glycans, showing glycogen accumulation in HCC-CC.

(E–F) Heatmaps of differentially expressed proteins (D) and transcripts (E) identified from unsupervised clustering of DBiT-GPT protein and transcriptome data, respectively.

(G) Spatial maps of liver zonation scores inferred from spatial transcriptomic annotations, showing the distributions of zone 1, zone 2, and zone 3 programs.

(H) Heatmap of transcripts dynamically regulated along pseudotime inferred from spatial transcriptomes, showing distinct transcript modules associated with the pseudotemporal transition.

(I) Left: Transcriptome-based pseudotime analysis from zone 3-like CH to HCC-CC–associated states. Right: other two representative transcripts dynamically changed along pseudotime.

(J) Significant canonical pathways identified by IPA (HCC-CC vs Zone 3-like CH).

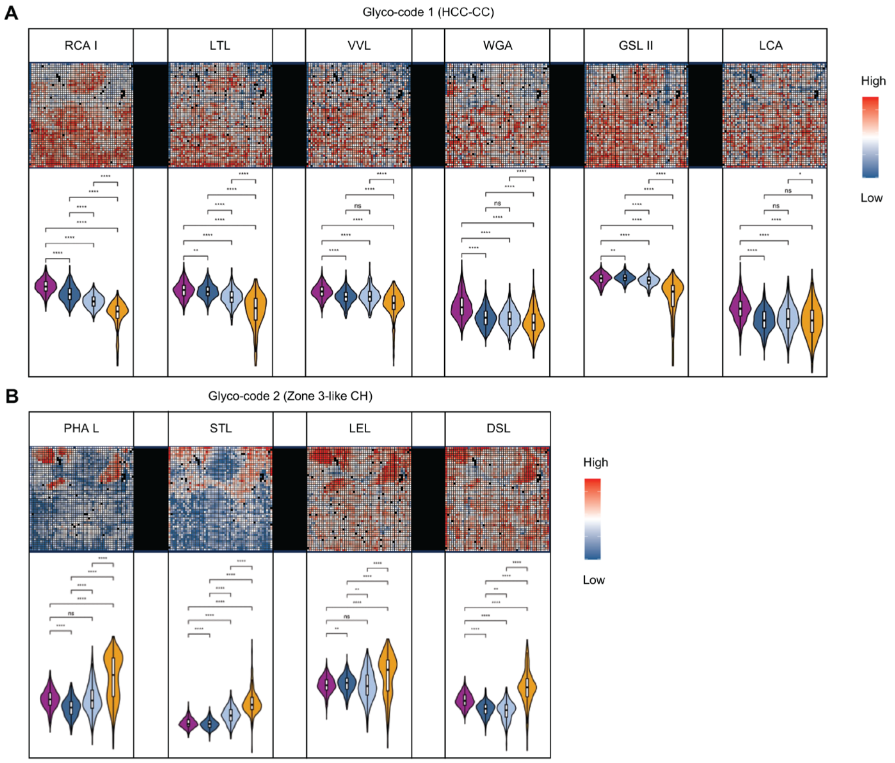


**Figure S8 Glyco-code and glycans expression of sample HCC-6T, related to Figure 3**

(A–B) Glyco-codes 1 and 2 identified in HCC-6T, corresponding to HCC-CC (A) and Zone 3-like CH (B), respectively. For each glyco-code, the upper panels show the spatial distribution of individual lectin-binding glycans, and the lower panels show the corresponding violin plots with statistical comparisons across clusters. Statistical significance across clusters was evaluated using a Kruskal-Wallis test followed by pairwise Wilcoxon rank-sum tests with Benjamini-Hochberg correction for multiple comparisons. Significance levels are indicated as ns, not significant; *, $P$adj < 0.05; **, $P$adj < 0.01; ***, $P$adj < 0.001; and ****, $P$adj < 0.0001.

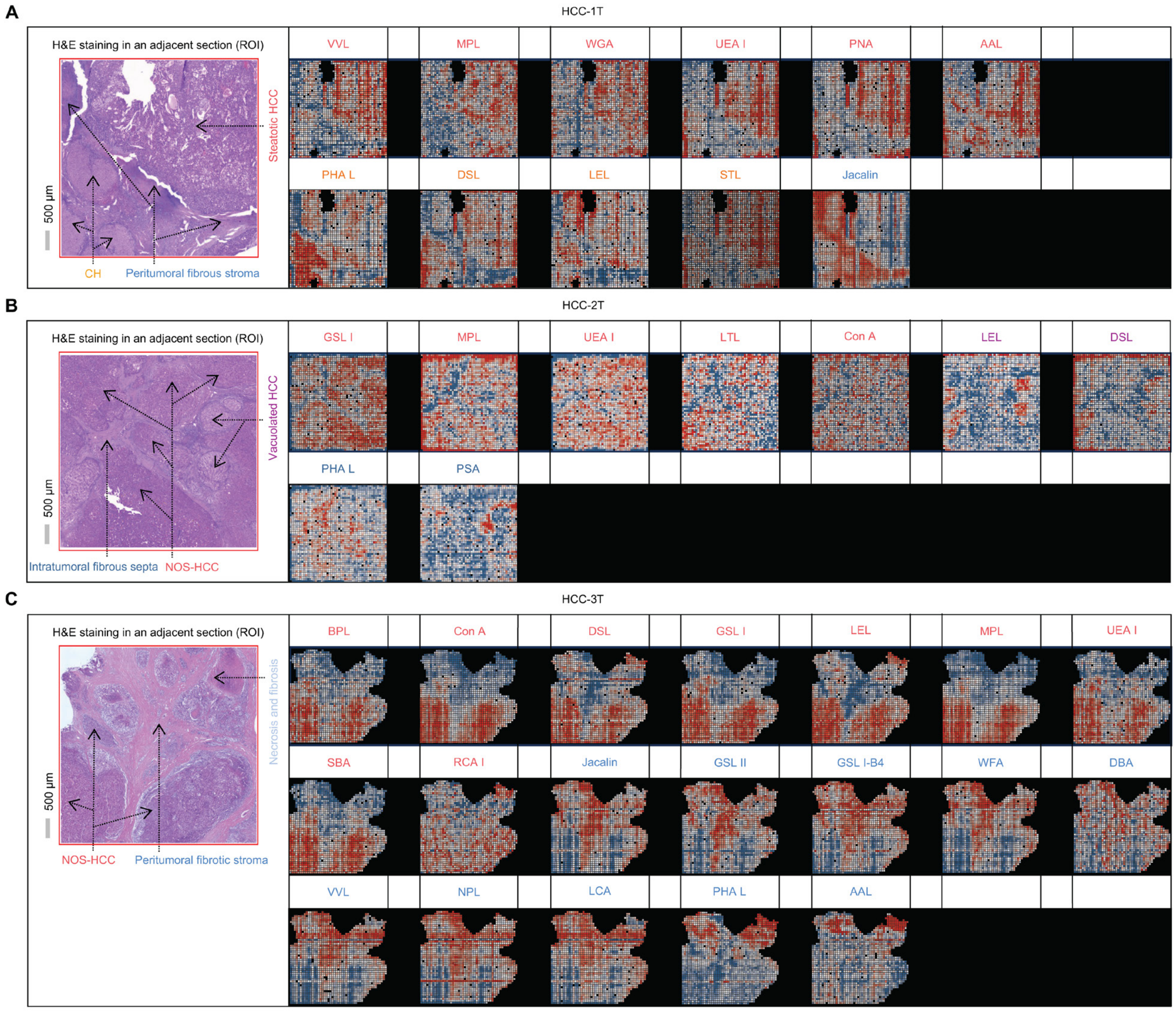


**Figure S9 Glycans expression of sample HCC-1T (A), HCC-2T (B), and HCC-3T (C), related to Figure 3**

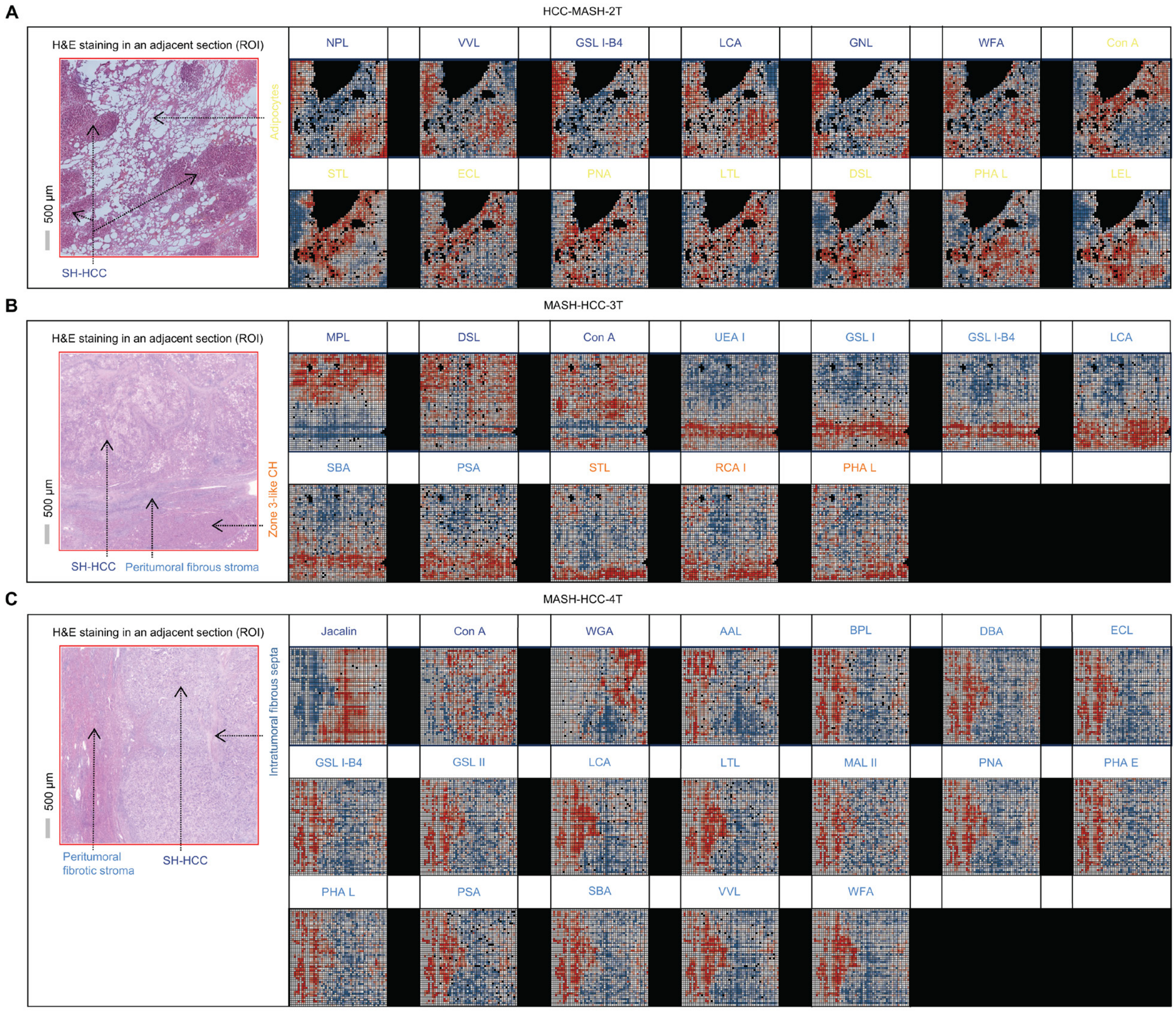


**Figure S10 Glycans expression of sample MASH-HCC-2T (A), MASH-HCC-3T (B), and MASH-HCC-4T (C), related to Figure 3**

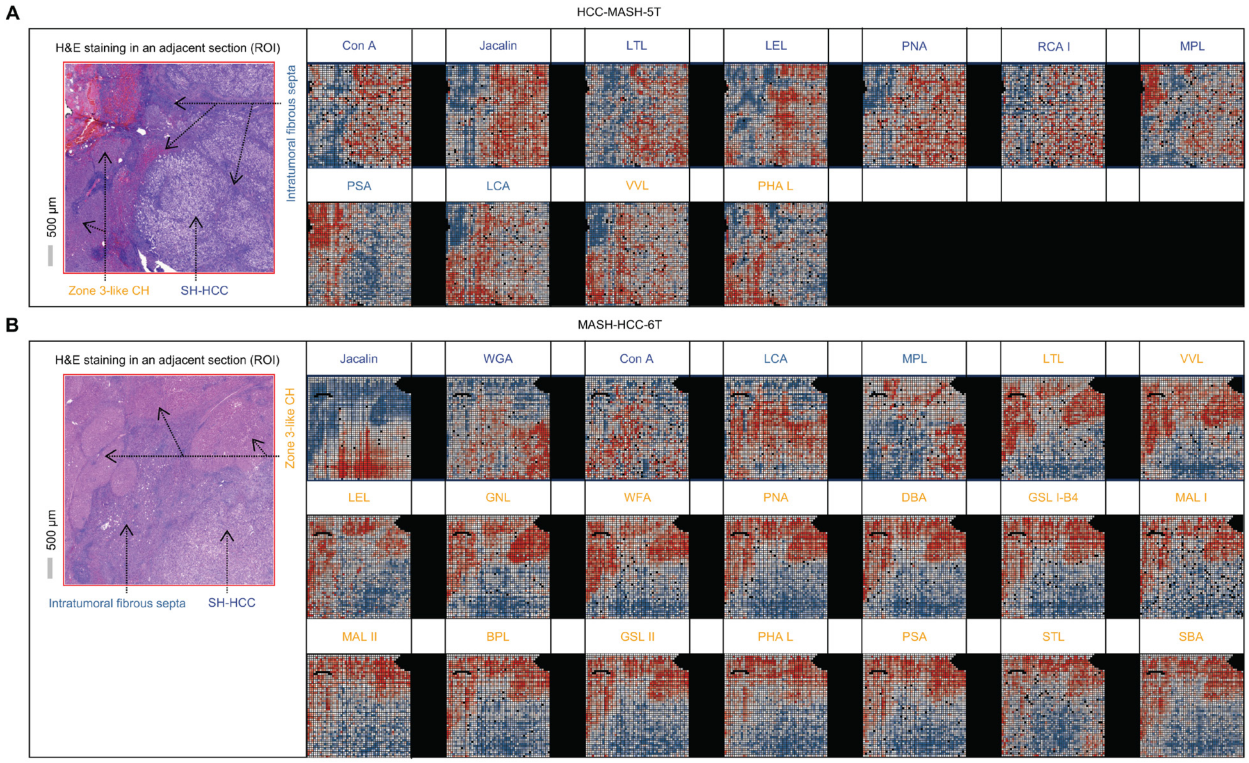


**Figure S11 Glycans expression of sample MASH-HCC-5T (A) and MASH-HCC-6T (B), related to Figure 3**

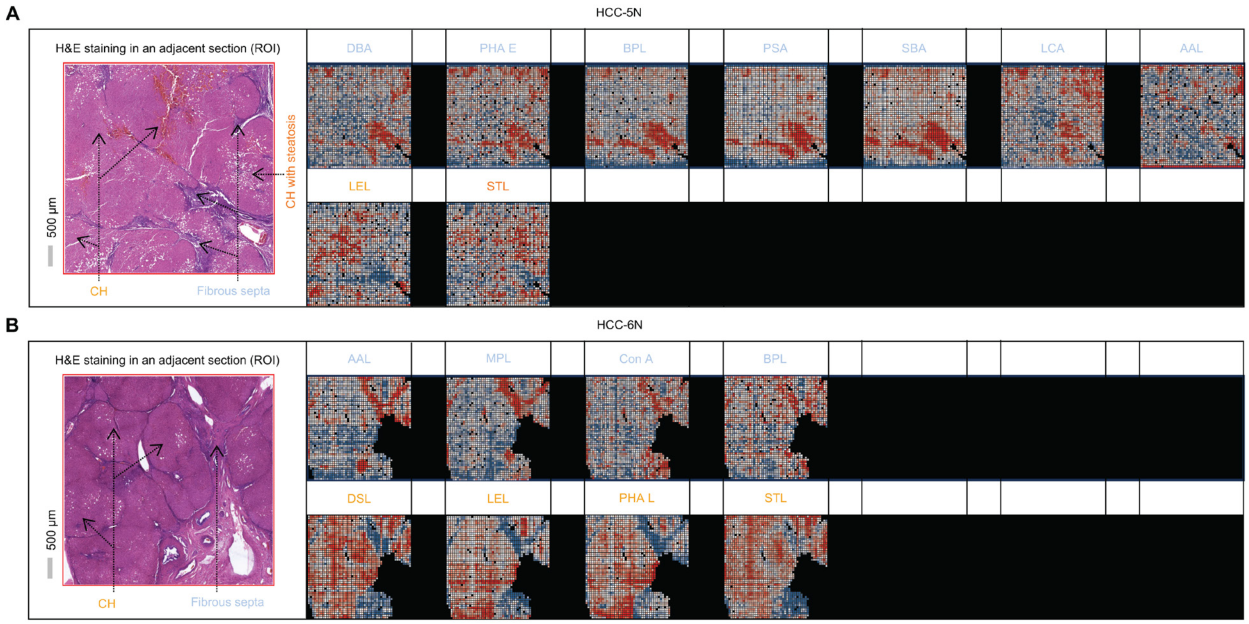


**Figure S12 Glycans expression of sample HCC-5N (A) and HCC-6N (B), related to Figure 3**

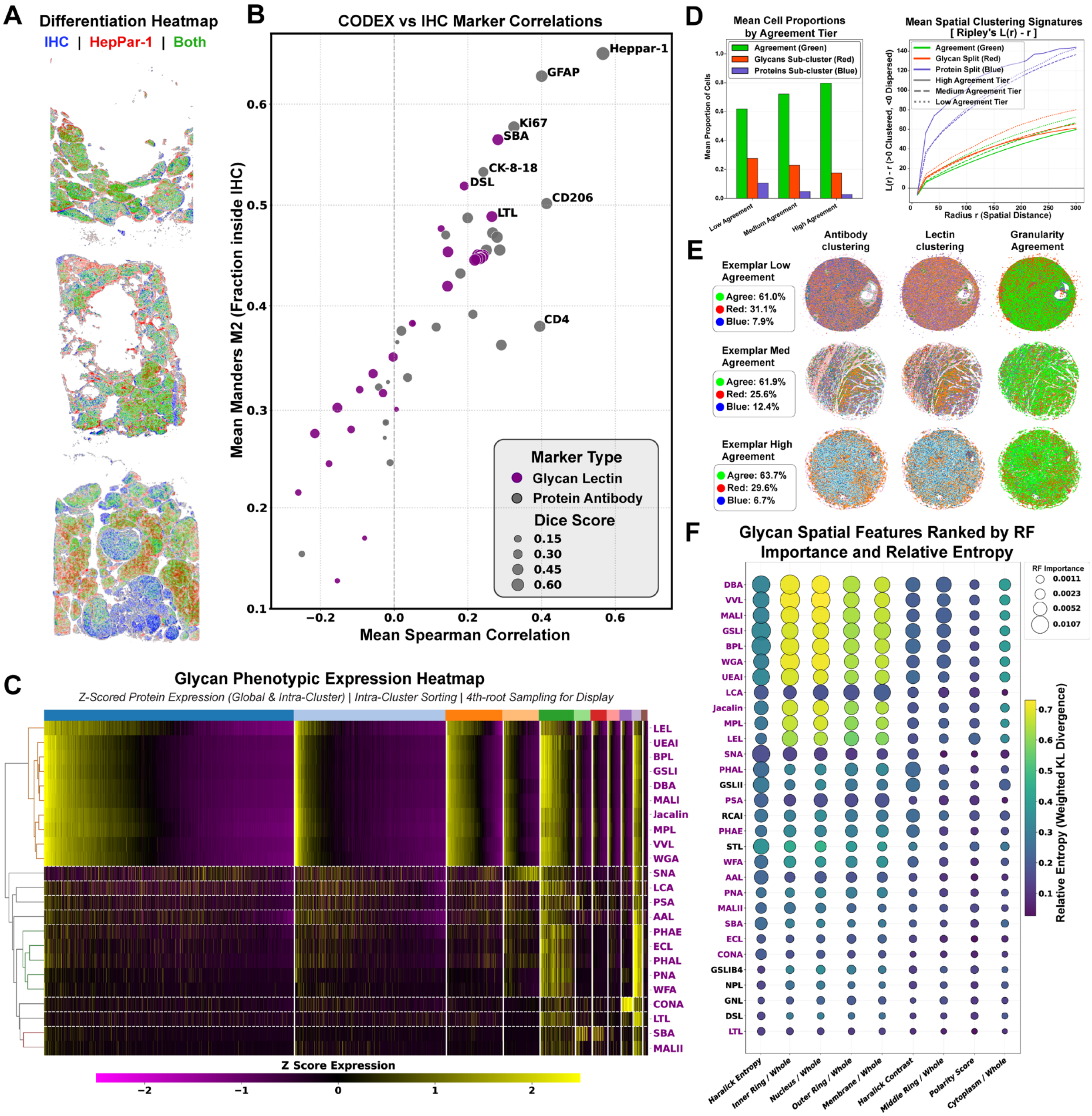

**Figure S13 CODEX-GP and IHC of MASH-HCC liver, related to Figure 4**
(A) Differentiation heatmap of adjacent sections for each biopsy, one stained with HepPar-1 IHC only, and the other stained with HepPar-1 mIF after lectin binding and non-specific blocking. Pixel colors were calculated according to the described methods in the main text. Generally, blue pixels correspond to positive IHC signal, red corresponds to positive CODEX signal, and green corresponds to concordance
(B) Correlation dot plot with selected top mIF correlators to HepPar-1 IHC signal, averaged across all three samples.
(C) Phenotypic heatmap clustered using both lectin and antibody markers.
(D) Examples of pooled granularity agreement tiers from all CODEX-GP samples and their spatial distribution characteristics. Note how the cell phenotypes where antibodies subcluster are distributed in smaller localized niches, especially in the high-agreement tier.

(E) Exemplar tissue cores clustered from CODEX-GP images using only antibodies or only lectins, color matched using a hybrid Hungarian and parent domination algorithm, with the granularity agreement maps for each agreement tier.
(F) Subcellular distribution features, their relative entropy, and their importance for select glycans. Manually selected glycans for clustering representations are highlighted in purple on the y axis.

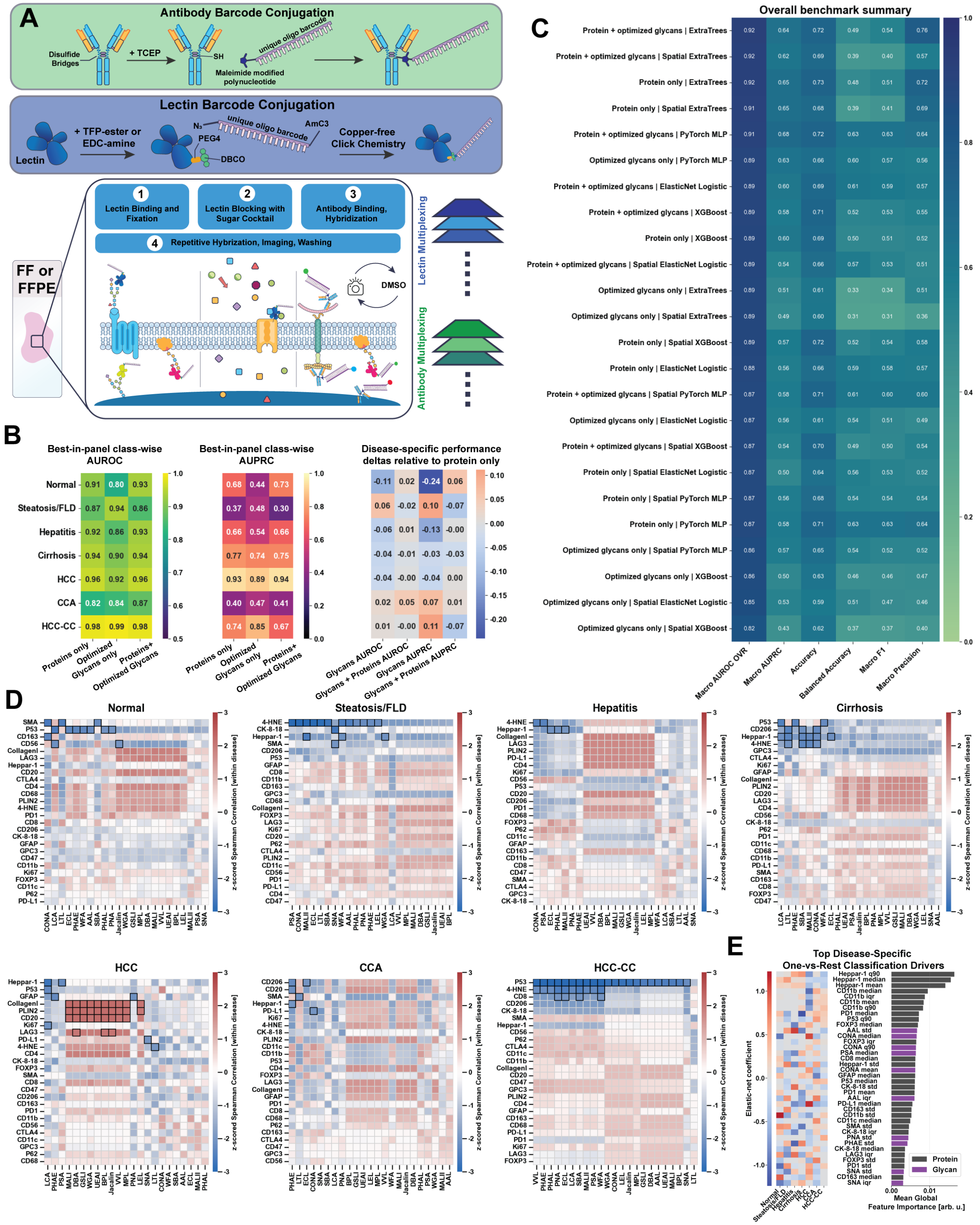

**Figure S14 CODEX-GP of multiple pathologies, related to Figures 4 and 5**

(A) Schematic workflow and chemical basis of CODEX-GP.

(B) Best-in-panel AUROC classification performance from multiple models for each disease using antibodies-only, lectins-only, and combined markers.

(C) Best-in-panel AUPRC classification performance from multiple models for each disease using antibodies-only, lectins-only, and combined markers.
(D) Select antibody-lectin z-scored Spearman correlations for each disease. Pairs with absolute Spearman correlations > 2 are highlighted with a bold bounding box. Disease-specific performance deltas relative to antibody-only performance is shown in the bottom right.
(E) Examples of pooled granularity agreement tiers from all CODEX-GP samples and their spatial distribution characteristics. Note how the cell phenotypes where antibodies subcluster are distributed in smaller localized niches, especially in the high-agreement tier.
(F) Top global feature importance from ExtraTrees classification across CV folds. HepPar-1 intensities scored the highest. Glycan motifs are red, proteins are blue.
(G) Disease specific one-vs-rest drivers of each disease classification. Positive coefficients indicate markers where their presence was helpful in affirmative disease classification.

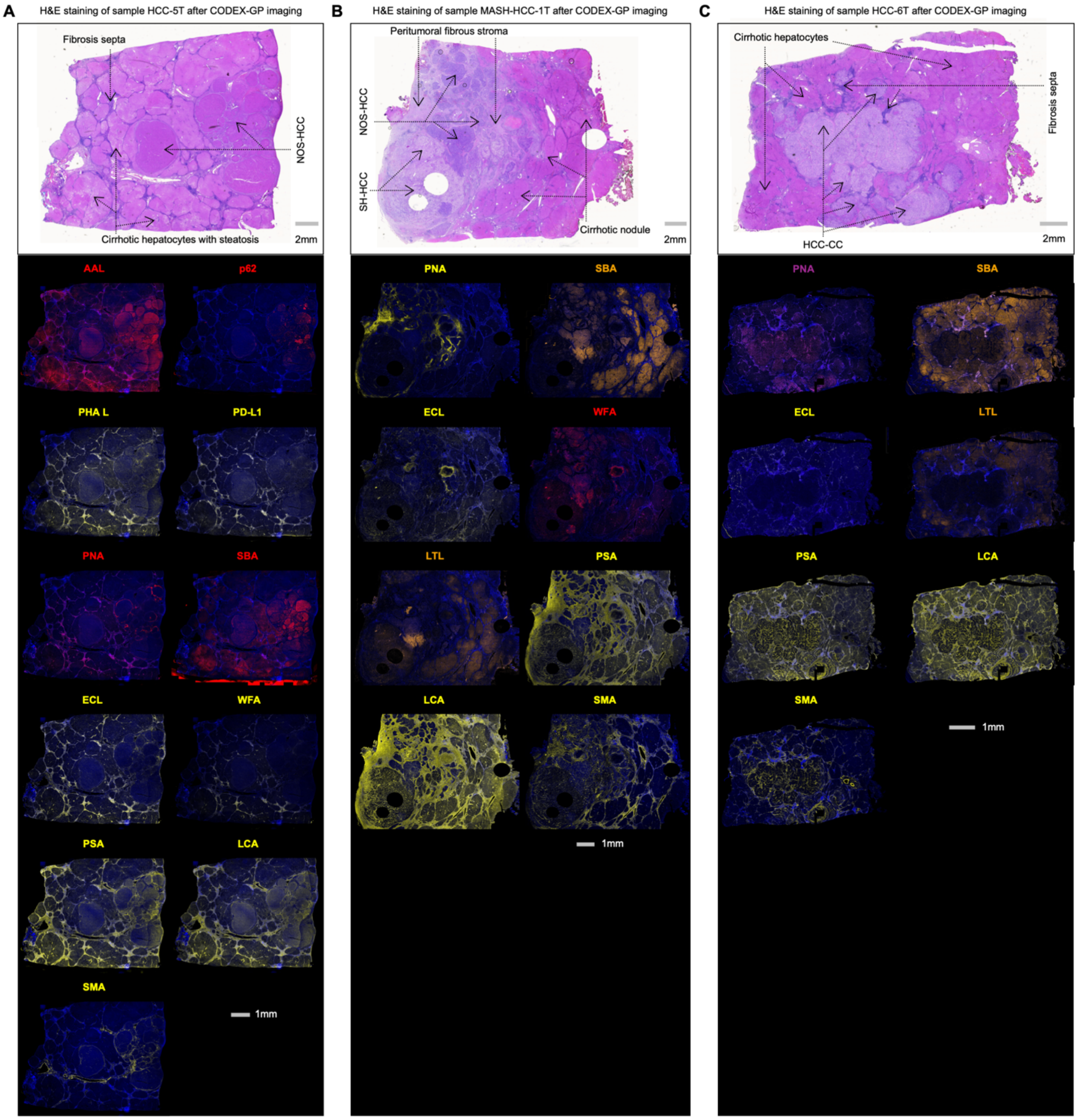


**Figure S15 CODEX-GP images of individual lectin- and protein-derived signals in samples HCC-5T, MASH-HCC-1T, and HCC-6T, related to Figure 4**

(A–C) CODEX-GP imaging of individual lectin- and protein-derived signals in samples HCC-5T (A), MASH-HCC-1T (B), and HCC-6T (C). The upper panels show H&E staining of the same tissue section after CODEX-GP imaging, with major histopathological regions annotated. The lower panels show selected lectin- and protein-derived signals with distinct spatial patterns across these pathological regions.
